# D-alanine aminotransferase (Dat) promotes *Staphylococcus aureus* colonization fitness on human nasal respiratory epithelium

**DOI:** 10.64898/2026.06.01.729472

**Authors:** Andrea I. Boyd, Katia A. Quintanilla, Isabel F. Escapa, Millie A. Lewis, Leah A. Kafer, Xi-Lei Zeng, Sarah E. Blutt, Carolyn B. Ibberson, Katherine P. Lemon

## Abstract

Nasal colonization by *Staphylococcus aureus* is an established risk factor for invasive infection, yet bacterial determinants promoting fitness on human nasal mucosa remain incompletely defined. To identify genes required for early colonization of human nasal respiratory epithelium, we colonized human nasal epithelial organoids differentiated at air-liquid interface (HNO-ALI) with a high-density transposon (Tn) library of the methicillin-resistant USA300 strain LAC. TnSeq analysis identified 165 genes that met our threshold for candidate colonization fitness factors. Among these, genes involved in D-alanine biosynthesis and use were enriched, including two encoding the enzymes that separately synthesize D-alanine in *S. aureus*: alanine racemase 1 (*alr1)* and D-alanine aminotransferase (*dat)*. Disruption of *dat* reduced colonization fitness in competition with the parental strain by ≥ 1,000 fold across 4 different strains from clonal complexes 8, 5, and 30. In competition with the parental strain during HNO-ALI colonization, a *dat*::Tn mutant was 34-fold less fit than an *alr1*::Tn mutant. Genetic complementation with single-copy *dat* expressed from its native operon promoter restored parental colonization levels. Supplementation with exogenous D-alanine or L-alanine also rescued the *dat*::Tn colonization defect, whereas D-glutamate did not, consistent with Dat primarily producing D-alanine on nasal mucosa. Complementation with *dat* under control of a putative 5’ intra-operon promoter substantially restored colonization but failed to support growth in chemically defined medium lacking L-alanine, suggesting a new layer of environment-specific regulation of *dat* transcription. Together, these findings demonstrate that Dat is a major source of D-alanine during colonization of human nasal mucosa and is required for *S. aureus* fitness in this environment.

**AUTHOR SUMMARY:** *Staphylococcus aureus* is the second leading cause of death due to bacterial infection globally, and nasal colonization is a major risk factor for invasive disease. Using a physiologically relevant, host-derived model of human nasal respiratory epithelium (HNO-ALI) and TnSeq, we identified 165 candidate genes contributing to *S. aureus* fitness during nasal mucosal colonization. We found that D-alanine aminotransferase (Dat) is the predominant source of D-alanine during nasal colonization, whereas alanine racemase (Alr1) predominates in rich medium, revealing an environment-specific hierarchy of D-alanine biosynthesis. Disruption of *dat* caused a > 1,000-fold defect in colonization in competition with the parental strain across multiple *S. aureus* clonal complexes, supporting a conserved role for *dat* in nasal colonization fitness. Additionally, we provide evidence that *dat* transcription from a previously cryptic promoter might be regulated by nasal mucosal conditions. Alr1 is proposed as an antimicrobial target in other bacterial pathogens; however, our data suggest that targeting Dat may be more effective for *S. aureus* nasal decolonization.

## INTRODUCTION

*Staphylococcus aureus* is the second leading cause of death due to bacterial infection globally, just behind *Mycobacterium tuberculosis* [1]. Human nasal colonization is a primary source for infection by *S. aureus*. Up to 80% of *S. aureus* infection isolates match the infected person’s nasal-colonizing strain [2–4]. Reducing nasal colonization can reduce subsequent *S. aureus* infections [5–8]. There is still no effective human vaccine to protect against *S. aureus* [9]. Without a vaccine, nasal decolonization with the topical antibiotic mupirocin has become a standard clinical approach to prevent infections in high-risk settings (e.g., preoperatively and in intensive care units) [10–13]. However, increased mupirocin use raises the risk of increasing antibiotic resistance rates [14, 15].

A key barrier to reducing the rate of *S. aureus* infections is identifying new methods to consistently eliminate or prevent nasal colonization. Transposon sequencing (TnSeq) screens have successfully identified genes required for *S. aureus* survival *in vitro* during antibiotic stress [16], nitric oxide exposure [17], intracellular infection of macrophages [18] and A549 cells [19], and *in vivo* during murine infections [19–23]. However, to date, there is only one prior *S. aureus* TnSeq screen of mucosal surface colonization and it is during murine vaginal colonization [24]. Although these model systems are excellent for identifying *S. aureus* genes important for specific treatment, host defenses, or systemic infection, we hypothesized that factors specific to the human nasal mucosal environment are not captured by these approaches. Supporting this, Bilici et al. recently identified a previously unknown role for biotin synthesis in nasal colonization, noting that biotin is abundant in cotton rat [25] and murine [26] plasma but limiting in human nasal secretions [25] and plasma.

During colonization, *S. aureus* is present throughout the nasal passages [27]. To identify *S. aureus* colonization fitness factors specific to the human nasal mucosal surface that coats the majority of the nasal passages, we used **h**uman **n**asal epithelial **o**rganoids differentiated at an **a**ir-**l**iquid **i**nterface (HNO-ALI). HNO-ALI accurately recapitulate human nasal respiratory epithelium with mucus-producing goblet cells, ciliated epithelial cells, and a thick mucus layer exposed to air on the apical surface [28–31]. HNO-ALI cultures are nontransformed and nonimmortalized, can be stored frozen, are stable over many passages, and support live bacterial colonization by human-nasal-associated species [31, 32]. These characteristics have enabled use of the same lines for experimentation over multiple years.

Here, we used HNO-ALI as a reductionist *ex vivo* model and TnSeq to screen a saturated*S. aureus* LAC transposon (Tn) library [17] and identified genes that contribute to early colonization on HNO-ALI. Among the strongest candidate colonization fitness genes were ones involved in D-alanine biosynthesis and use. While humans lack D-alanine, in bacteria, D-alanine is essential for peptidoglycan crosslinking [33] and for D-alanylation of gram-positive teichoic acids, which modulates cell wall charge and resistance to cationic antimicrobial peptides [34, 35].

*S. aureus* produces D-alanine via two ways. The racemase Alr1 interconverts L-alanine and D-alanine and is the primary source of D-alanine in culture medium [36]. In addition, the aminotransferase Dat interconverts D-glutamate and pyruvate with D-alanine and 2-oxoglutarate. D-alanine synthesis has been targeted in *S. aureus* [37] and other bacterial species primarily by inhibiting Alr1 [38–40]. Although *S. aureus* can generate D-alanine using either Alr1 or Dat, Panda et al. show Dat contributes only minimally to D-alanine pools in wild-type *S. aureus* growing in a chemically defined medium, with Dat thought to primarily produce D-glutamate [36]. *S. aureus* is also annotated as having a second alanine racemase (*alr2*) but *alr2* alone is insufficient to enable growth in an *alr1 dat* double mutant in the absence of exogenous D-alanine [36, 41]. An *alr1 alr2 dat* triple mutant also has substantially reduced virulence in mice [41].

The HNO-ALI model revealed a predominant role for *dat* during *S. aureus* colonization of a human nasal mucosal surface. Tn disruption of *dat* resulted in a more profound defect during monocolonization on HNO-ALI (∼7000-fold decrease relative to the parental strain level) than did disruption of *alr1* (∼54-fold decrease relative to the parental strain level). When competed directly against the parent in cocolonization of HNO-ALI, the *dat*::Tn mutant consistently had a larger fitness defect than did *alr1*::Tn. Genetic complementation restored each mutant to parental levels, including new evidence for a possible intra-operon promoter immediately 5’ of *dat* that was functional on HNO-ALI but not in a chemically defined medium lacking L-alanine. Moreover, exogenous D-alanine but not D-glutamate was sufficient to rescue the *dat*::Tn colonization defect, indicating that production of D-alanine is a primary role for Dat during human nasal mucosal colonization. Additionally, *dat* was an important colonization fitness factor in *S. aureus* strains from clonal complexes (CC) 8, 5, and 30. Overall, this study points to the predominance of Dat over Alr1 for biosynthesis of the essential amino acid D-alanine during *S. aureus* colonization of human nasal mucosa, and suggests that targeting Dat, rather than Alr1, could be a preferred strategy for *S. aureus* nasal decolonization.

## RESULTS

### TnSeq identifies *S. aureus* genes for the biosynthesis and use of D-alanine as candidate fitness factors for colonization of human nasal mucosa

*S. aureus* nasal colonization is an important area of research because of its clinical importance as a key risk factor for *S. aureus* infection. Studies using animal models reveal roles of the immune system in *S. aureus* nasal colonization [42]. Studies using cancer-derived cells lines such as RPMI 2650 cells have helped identify *S. aureus* adhesion factors [43], and interactions with other members of the human nasal microbiota [44, 45]. However, we hypothesized that there are as-yet-unidentified factors unique to the human nasal mucosal surface that influence *S. aureus* colonization. To address this, we used the reductionist *ex vivo* model system of HNO-ALI, which contain all epithelial cell types and have an apical mucus layer in direct contact with air. We then colonized HNO-ALI with the methicillin-resistant *S. aureus* LAC Tn library developed by Grosser and colleagues [17]. This is a saturated library with 77,161 independent insertions in the LAC genome [46]. In order to capture *S. aureus* genes important for early colonization rather than for infection, we colonized HNO-ALI for 8 h at the human nasal physiological temperature of 34 °C [47], since *S. aureus* behavior shifts towards virulence at 37 °C and above [48]. The Baylor College of Medicine 3D Organoid Core recently revised their protocol for generating HNO-ALI, which has improved timely and consistent production. However, with these new HNO-ALI, *S. aureus* JE2 (a derivative of LAC) colonization induced cytotoxicity by 24 h in an *hla*-dependent manner (**Fig. S1A**) but its parent strain LAC was well tolerated at 8 h (**Fig. S1B**). This 24-hour damage phenotype is consistent with *S. aureus*’*s* known *hla*-dependent damage of polarized cancer-derived cell lines [49]. Therefore, we used an 8-hour timepoint to identify *S. aureus* genes important for fitness early in human nasal mucosal colonization, before any *hla*-dependent damage occurred.

We detected Tn insertions in 2,505 genes in the input TnSeq library, and in 2,434 genes after 8 hours (h) of HNO-ALI colonization (**Table S1A**). To identify genes with a disproportionately underrepresented number of Tn insertions after colonization compared to the input library (indicating importance for fitness on HNO-ALI), we first normalized both the input and output to a simulated library of all expected Tn insertions [18, 22, 50–52]. (There were 388 genes with underrepresented Tn insertions in the input library and 462 in the output compared to the simulated library.) We then plotted the log2 fold change (log2FC) between observed and simulated insertion sites in the input library on the y-axis and in the output after HNO-ALI colonization on the x-axis (**Fig. 1A**). To identify candidate fitness genes, we set a threshold for log2FC ≤ −1.5 for Tn presence after colonization compared to input. 165 genes met this threshold (light blue or red in **Fig. 1A**).

**Figure 1.**
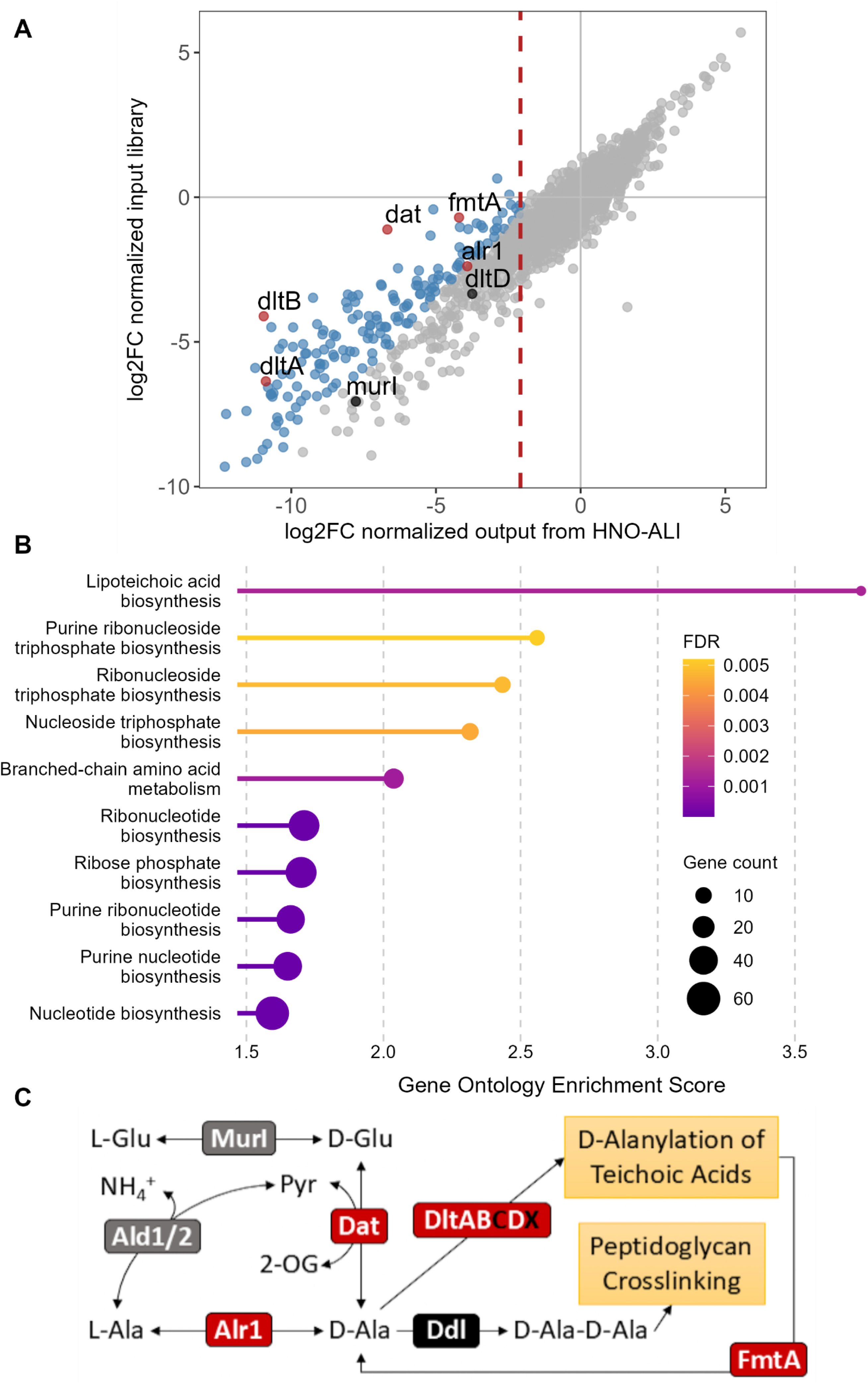
>*S. aureus* genes encoding proteins involved in cell wall integrity and charge are candidate colonization fitness factors on a human nasal mucosal surface. **(A)** Scatter plot of gene fitness normalized to a simulated library of all expected Tn insertions in *S. aureus* LAC after 8 h on HNO-ALI (x-axis) at 34 °C compared to the normalized input Tn library (y-axis); *n* = 2 independent experiments in HNO919 (**Table S1** for full data). Tn insertions in all genes to the left of the dotted red line were significantly underrepresented compared to the simulated library after HNO-ALI colonization with a threshold of log2FC ≤ −2, and a *p*-value of < 0.05. Light blue indicates genes within this subset for which Tn insertions were further underrepresented after HNO-ALI colonization relative to the input library (Δlog₂FC ≤ −1.5, defined as log2FC output - log2FC input). Genes involved in D-alanine synthesis or utilization are highlighted in red if they meet the Δlog₂FC threshold and in black if they do not. Gray indicates all other genes. (**B**) Enrichment analysis using STRING and Biological Processes (Gene Ontology) identified pathways enriched for genes contributing to fitness during HNO-ALI colonization. FDR and gene count are indicated on the panel. (**C**) Schematic of the *S. aureus* D-alanine synthesis pathway and cell-wall uses. Protein names are colored red if Tn insertions in the corresponding genes were underrepresented in the output. (Tn insertions in *ddl*, *dltC,* and *dltX* were considered absent in the input library and *murI* just met the threshold for “present” in the input library.)

To identify pathways enriched for candidate fitness genes, we performed an enrichment analysis using STRING with the Biological Processes pathways from Gene Ontology [53–55] (**Fig. 1B** and **Table S1B**). The most highly enriched pathway was “lipoteichoic acid biosynthetic processes”. The fifth most enriched was “branched chain-amino acid and metabolic processes”. The remaining eight of the top ten enriched pathways were all involved in synthesis of purines and/or nucleotides/sides.

A number of these candidate fitness genes corroborate data from prior screens for *S. aureus* fitness factors in a host environment, supporting HNO-ALI as a model (**Table S1A**). A large portion of genes with significantly fewer Tn insertions have roles in nutrient acquisition and metabolism. For example, *cntE* and *ccpA* were identified as important for *S. aureus* fitness during nasal colonization and play a role in fitness in a murine model of vaginal colonization [24]. *cntE* encodes an MFS transporter that exports staphylopine, a metallophore that scavenges a variety of metals countering host nutritional immunity [56]. *ccpA* (carbon catabolite protein A) is a major regulator of *S. aureus* transcription in response to the availability of glucose and other carbon sources [57, 58]. Purine synthesis is well-established to contribute to *S. aureus* infection [20, 59–62]. We find many genes related to purine synthesis, including *purB, purD, purE*, and *purH*, are required for *de novo* purine biosynthesis and have a role in fitness during murine vaginal colonization [24], as well as in human nasal mucosal colonization (**Fig 1B**). Detection of these genes as candidate fitness factors for *S. aureus* early colonization of HNO-ALI corroborates data from other models for understanding of *S. aureus* colonization of host mucosal surfaces.

Three of the top 165 genes from our TnSeq screen are involved in D-alanine synthesis or use (**Fig. 1A**): *alr1* (alanine racemase 1), *dat* (D-alanine aminotransferase) and *fmtA* (teichoic acid D-alanine esterase). This is consistent with D-alanine’s role as an essential amino acid for cell wall crosslinking in bacteria and for cell-surface charge modulation in gram-positive bacteria. FmtA cleaves D-alanine from lipoteichoic acids (LTA) to provide D-alanine for attachment to wall teichoic acids (WTA) [63], modulating cell-surface charge. A double *alr1 dat* mutant is auxotrophic for D- alanine [36, 41]. Alr1, a racemase, reversibly interconverts L-alanine and D-alanine while Dat, an aminotransferase, reversibly interconverts pyruvate and D-glutamate with 2-oxoglutarate and D-alanine (**Fig. 1C**). In culture media, Alr1 is the primary producer of D-alanine [36]. In contrast to its expected preeminence, *dat*::Tn had much greater underrepresentation than *alr1*::Tn in our TnSeq screen during early colonization of HNO-ALI (**Fig. 1A**).

Other genes involved in D-alanine usage were also either disproportionately underrepresented after colonization (e.g., *dltAB*) or were absent in the input Tn library (*ddl, dltC, dltX*) (**Fig. 1C**). We considered genes with < 10 Tn insertions in each of our independent screening experiments as absent in the input library (highlighted in black in **Fig. 1C**). The function of each gene in the D-alanine teichoic acid operon (*dltABCDX*) is still under investigation; however, the current model supports that *dltA* adenylates D-alanine for transfer to carrier protein *dltC*, and *dltC* transfers D-alanine to lipoteichoic acid via a conformational change with *dltB* [64]. *murI* is a glutamate racemase that interconverts L-glutamate and D-glutamate [65, 66], whereas *ddl* is required for crosslinking cell wall peptidoglycan and is essential in *S. aureus* under a variety of growth conditions [67].

### Disruption of *dat* results in a greater reduction in *S. aureus* fitness during colonization of a human nasal mucosal surface than does disruption of *alr1*

Based on their underrepresentation in the TnSeq screen, we hypothesized that *alr1*::Tn, *dat*::Tn, and *fmtA*::Tn mutants are less fit than the parental *S. aureus* strain during HNO-ALI colonization. To better assess the importance of these genes during colonization, we performed the following experiments in *S. aureus* JE2 *hla*::Tn(spc) strain (KPL4530), since an *hla*-deficient background extends the colonization time on HNO-ALI (**Fig. S1A**). We transduced each Tn(erm) mutant of interest from the NTML into this parental background and, unless stated otherwise, all subsequent mutants referred to here are double mutants with *hla*::Tn(spc).

We determined the fitness of each mutant compared to the parent strain by cocolonizing HNO-ALI with equivalent amounts (1x105 CFUs) of each strain with the parent for 24 h at 34 °C with recovered CFUs per HNO-ALI as the output. (As controls, we also monocolonized HNO-ALI with each strain, available at https://klemonlab.github.io/DatCol_Manuscript/.) The parental *S. aureus* strain yielded 107-to-108 CFUs/HNO-ALI in all cocolonization conditions and outcompeted each of the mutants. On average, we recovered approximately 1,502-fold fewer CFUs *dat*::Tn, 34.1-fold fewer *alr1*::Tn, and 5.08-fold fewer *fmtA*::Tn than the parent after 24 h of cocolonization (**Fig. 2A**). The competition index (CI) was calculated as (mutant output / parent output) / (mutant input / parent input) to quantify reduced fitness compared to the parent *S. aureus*. Each mutant had a CI significantly less than 1, where 1 indicates fitness equal to the parental strain (*p* < 0.05). The model-predicted mean competition index was 0.00077 for *dat*::Tn, 0.026 for *alr1*::Tn, and 0.12 for *fmtA*::Tn (**Fig. 2B** and **Table S2A**). Although *fmtA*::Tn had a statistically significant colonization fitness defect, it was milder than that of the other two mutants. The *dat*::Tn CI was significantly lower than that of *alr1*::Tn by 34-fold (*p* = 0.002). These data demonstrate that the *dat*::Tn mutant was significantly less fit than the *alr1*::Tn mutant during nasal colonization on HNO-ALI.

**Figure 2.**
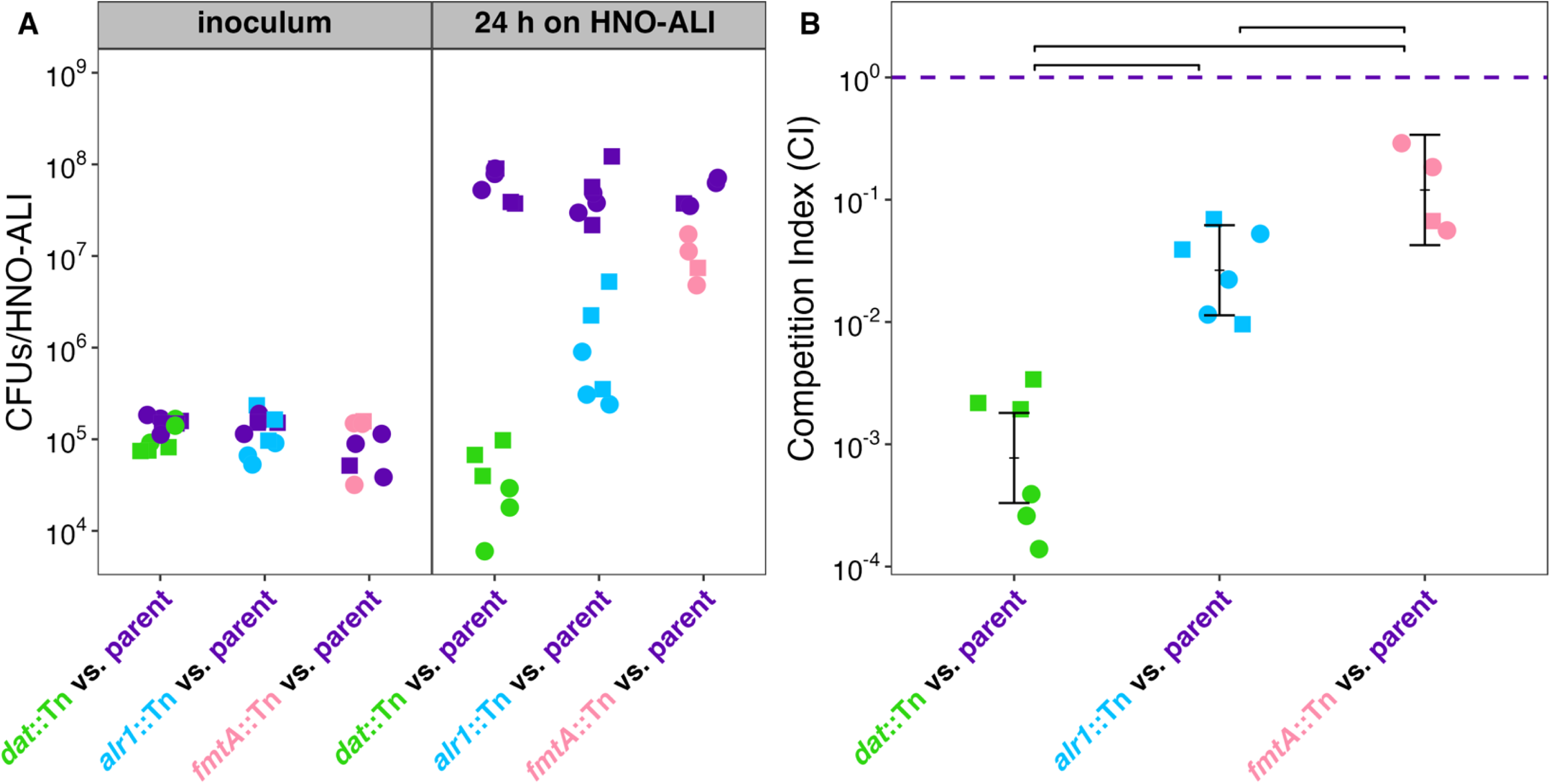
Genes involved in the synthesis and use of D-alanine are required for full *S. aureus* fitness on a human nasal mucosal surface. (**A**) The parental strain (purple) outcompeted an isogenic mutant with a Tn(erm) disruption in either *dat* (green), *alr1* (blue), or *fmtA* (pink), respectively. HNO-ALI were cocolonized with 10^5^ CFUS/HNO-ALI each of the parental *S. aureus* strain KPL4530 (JE2 with *hla*::Tn(spc)) and one mutant of interest for 24 h at 34 °C in *n* = 3 independent experiments in each of 2 HNO lines (HNO204, circles and HNO918, squares). Competition was done only once in HNO918 for *fmtA*::Tn. (**B**) Competition indices (CI = (mutant output / parent output) / (mutant input / parent input)) of data from panel A show decreased fitness for the three mutants compared to the isogenic parental strain. Log-transformed CI were modeled with mutant as a fixed effect and experimental date as a random effect. All pairwise contrasts were performed and adjusted for multiple comparisons using the Holm method. Horizontal brackets represent statistically significant comparisons *p* < 0.05. Vertical brackets represent model-predicted mean values and confidence intervals (± 2 × SEM), back-transformed to the original response scale. The purple dashed line indicates a CI value of 1, representing equal fitness between the mutant and the parental strain, and serves as the reference value for the one-sample t-tests performed for each strain, all of which were statistically significant (*p* < 0.05). Statistical analysis is in **Table S2A**.

### Genetic complementation of *dat*::Tn restores *in vitro* growth in alanine-deficient chemically defined medium

Because of its evident contribution to *S. aureus* fitness, we focused on investigating why *dat* is so critical during nasal colonization. We first evaluated whether the colonization defect in the *dat*::Tn mutant was due solely to the Tn insertion in the *dat* gene via single copy complementation with pLL39 integrated at the Φ11 *attB* site [68]. The *dat* gene is the second member of a two-gene operon, preceded by *pepV* (aka *sapep*) [36], which encodes a secreted dipeptidase [69, 70]. Panda et al. show that the promoter 5’ of *pepV* (P*pepV*) followed by either the ribosomal binding site of *pepV* or of *dat* plus the *dat* gene is sufficient to complement a *dat* deletion [36]. We therefore placed P*pepVdat* at the Φ11 *attB* site in both the *dat*::Tn and parent strains. In addition, we used SAPPHIRE [71] to predict potential promoters in this operon region (**Table S3**) to hypothesize the presence of a cryptic promoter internal to the operon in the region immediately 5’ of *dat*. To test this, we placed a fragment consisting of a 371 bp region 5’ of the start codon in *dat* plus the *dat* gene (P*datdat*) at the Φ11 *attB* site in both the *dat*::Tn and parent strains (**Fig. S2A**). We whole genome sequenced *dat*::Tn P*datdat* and found P*datdat* was inserted at the *attB* site in the opposite direction of all adjacent genes (**Fig. S2B**) supporting *dat* transcription from this region is due to the hypothesized promoter. We also generated a construct to complement *alr1* with its native promoter.

We hypothesized that the mutant and complemented strains would have equivalent growth curves in nutritionally replete medium. To test this, we grew each strain separately in tryptic soy broth (TSB) and a MOPS-based chemically defined medium supplemented with glucose and all 20 proteogenic L-amino acids (CDM20) [72] for 20 h (**Fig. S3A-C**). All strains had equivalent growth curves in CDM20. The Δ*alr1* mutant had a slightly reduced area under the curve (AUC) in TSB compared to the parent strain KPL4530 and compared to the parental strain with the empty vector pLL39 (EV) (<12% less AUC) (**Table S2B**).

An *S. aureus alr1 dat* double mutant is an auxotroph for D-alanine [36, 41]. Therefore, in a *dat*::Tn mutant, generation of D-alanine is dependent on *alr1* (via Alr1 conversion of L-to D-alanine (**Fig. 1C**)). Consequently, a *dat*::Tn mutant should be incapable of growth in alanine-deficient CDM (CDM19-ala), which lacks the needed L-alanine substrate for Alr1. Furthermore, we hypothesized that this defect would be rescued by single-copy complementation with either P*pepVdat* or P*datdat* restoring the ability to convert D-glutamate to D-alanine. As expected, the *dat*::Tn mutant failed to grow in CDM19-ala (**Fig. 3A**). Integration of P*pepVdat* in single copy at the Φ11 *attB* site in both *dat*::Tn and the parental strain resulted in greater growth than that of the parental strain with the empty vector pLL39 (EV) (**Fig. 3A**). This demonstrates that complementation with single-copy P*pepVdat* is sufficient to rescue the growth defect caused by a *dat*::Tn mutation in the absence of exogenous L-alanine, consistent with previous literature [36]. It also suggests that, in the presence of intact *alr1* and the absence of L-alanine, increasing the level of Dat above what is produced from its wild-type chromosomal location by adding P*pepVdat* to the parental strain results in increased growth. This is most likely due to increased production of D-alanine, which can be used for cell wall integrity and/or conversion to L-alanine for protein synthesis. Furthermore, the increased growth of the *dat*::Tn P*pepVdat* compared to the parental strain implies that placing P*pepVdat* at the Φ11 *attB* site and/or placing P*pepV* directly 5’ of *dat* increases the levels of Dat in the cell compared to when *dat* is in its wild-type chromosomal position. In contrast to the complementation of *dat*::Tn by P*pepVdat*, integration of P*datdat* at the Φ11 *attB* site failed to rescue growth of the *dat*::Tn and had no effect on growth of the parental strain (**Fig. 3A**).

**Figure 3.**
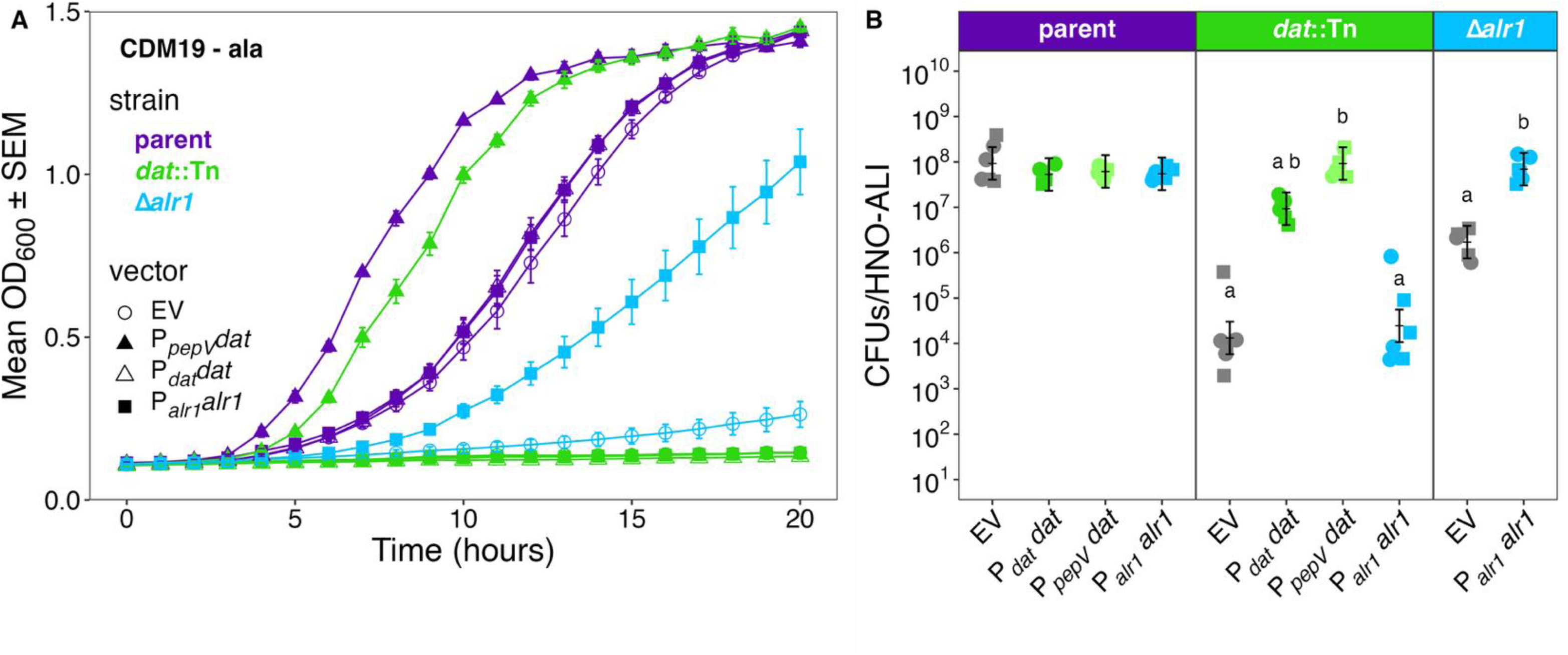
Genetic complementation of both the *S. aureus dat*::Tn and Δ*alr1* mutant restores parental-level colonization on a human nasal mucosal surface. The parental *S. aureus* KPL4530 (JE2 *hla*::Tn(spc)) (purple) and isogenic *dat*::Tn(erm) (green) and Δ*alr1* (blue) mutants each carried either the empty vector (EV) pLL39 (empty circles), P*_pepV_dat* (filled triangles), P*_dat_dat* (empty triangles), or P*_alr1_alr1* (filled squares), inserted at the Φ11 *attB* site. Growth of each strain with no vector is in **Fig. S3C**. (**A**) Genetic complementation restored growth in liquid CDM minus alanine (CDM19-ala) fully for *dat*::Tn P*_pepV_dat,* partially for Δ*alr1* P*_alr1_alr1*, and not at all for *dat*::Tn P*_dat_dat.* All strains were grown in CDM19-ala for 20 h in *n* = 3 independent experiments. Each data point is the average OD600 ± SEM of all 3 experiments at that time point. (**B**) Strains indicated on the x-axis were colonized on HNO-ALI for 24 h in *n* = 3 independent experiments in each of 2 HNO lines (HNO204, circles and HNO918, squares). Log-transformed CFUs/HNO-ALI between groups were compared using an LMM to determine statistical significance with background strain and vector as fixed effects and experimental date as a random effect. Pairwise contrasts were performed to compare all conditions to the parent strain carrying the same vector (labeled “a” if Holm-adjusted *p* < 0.05) and to compare each vector-bearing mutant to its isogenic empty-vector mutant strain (labeled “b” if Holm-adjusted *p* < 0.05). Vertical brackets represent model-predicted mean values and confidence intervals (± 2 × SEM), back-transformed to the original response scale. Statistical analysis is in **Tables S2B-C**.

### In the absence of exogenous L-alanine, a*lr1* is required for *in vitro* growth in CDM, implying a key role for *alr1* in supporting endogenous levels of L-alanine for protein synthesis under these conditions

In the absence of exogenous L-alanine, an Δ*alr1* mutant is dependent on *dat* for D-alanine production and on other pathways (e.g., *ald1*/*2*) for production of L-alanine for protein synthesis. The failure of the Δ*alr1* mutant to grow more than minimally in CDM19-ala (**Fig. 3A**), implies two possibilities, either 1) Dat alone produces an insufficient level of D-alanine under these growth conditions, which is less likely since in the absence of exogenous L-alanine the parental strain grows, or 2) in the absence of both exogenous L-alanine and Alr1, there is insufficient endogenous production of L-alanine via other routes to support protein synthesis and growth. Complementation of Δ*alr1* with single-copy P*alr1alr1* at Φ11 *attB* site partially restored growth. These data imply both that Alr1 is required for growth in the absence of exogenous L-alanine and that placing P*alr1alr1* at the Φ11 *attB* site results in lower levels of Alr1 protein than is produced from the parental chromosomal position. Our focus here is on nasal mucosal colonization; therefore, these possibilities are left for future investigation.

### Genetic complementation of *dat*::Tn or Δ*alr1* during *ex vivo* colonization on HNO-ALI point to influences of the nasal mucosal environmental conditions

To test for genetic complementation *ex vivo* on a nasal mucosal surface, we colonized HNO-ALI for 24 h with the parental *S. aureus* strain separately carrying each complementation construct (EV, P*pepVdat*, P*datdat,* or P*alr1alr1*), *dat*::Tn and Δ*alr1* carrying the EV, and each mutant with its respective complementation constructs. We also gave the *dat*::Tn strain an extra copy of *alr1* to determine if that would improve its colonization (**Fig. 3B**). HNO-ALI colonization by the parental *S. aureus* strain was unaffected by any of the 4 vector constructs (**Fig. 3B**). We recovered a mean CFUs/HNO-ALI of 107-to-108 parental *S. aureus* with the EV, P*pepVdat,* P*datdat,* or P*alr1alr1.* We then compared each mutant strain to its parental counterpart with its matched vector (for example, parent P*pepVdat* compared to *dat*::Tn P*pepVdat*; labeled as ‘a’ if significant) and each complemented mutant to its matched mutant with the EV (pLL39, labeled as ‘b’ if significant).

The *dat*::Tn EV strain had a 7,005-fold monocolonization defect compared to the parent EV strain. Single-copy P*pepVdat* in *dat*::Tn was sufficient to completely restore a parental level of nasal colonization (with 6,954-fold more CFUs/HNO-ALI than for *dat*::Tn EV) (**Fig. 3B**). In contrast to the lack of complementation during *in vitro* growth in CDM19-ala, we recovered 701-fold more *dat*::Tn P*datdat* than *dat*::Tn EV, providing complementation to near-parental levels with only 5.7-fold fewer than the parent P*datdat* strain. This substantial rescue of the *dat*::Tn colonization defect on HNO-ALI indicates that there is, in fact, a promoter within the 371 bp 5’ of the *dat* start codon, which is both internal to the bicistronic operon and overlaps with the coding sequence of *pepV* (the first gene in the operon). The near parental-level complementation of *dat*::Tn on HNO-ALI compared to the lack of any effect in CDM19-ala, points to regulation of the P*dat* in this region by environmental conditions. It also points to a path for future investigation of these conditions using a P*dat*-driven reporter construct.

Giving *dat*::Tn an extra copy of P*alr1alr1* did not have a statistically significantly effect on colonization levels (with only a 1.85-fold increase in *dat*::Tn *alr1* CFUs/HNO-ALI compared to *dat*::Tn EV) (**Fig. 3B**). Therefore, under the environmental conditions on HNO-ALI, an extra copy of *alr1* is insufficient to improve *dat*::Tn colonization. This implies that native production of Alr1 on HNO-ALI is sufficient to use the available exogenous L-alanine (and that there is unlikely to be excess L-alanine availability on HNO-ALI).

The Δ*alr1* EV displayed a modest colonization defect compared to parent EV with 54-fold fewer CFUs/HNO-ALI, pointing to a role for *alr1*-dependent production of L-alanine (for protein synthesis) and/or D-alanine (for cell-wall integrity) during nasal mucosal colonization. In contrast to the partial complementation during *in vitro* growth on CDM19-ala, during *ex vivo* colonization of HNO-ALI, single-copy P*alr1alr1* fully restored nasal mucosal colonization at 24 h on HNO-ALI to parental levels (**Fig. 3B**). We expect this difference to be due to some availability of L-alanine in HNO-ALI mucus, consistent with the known detection of alanine in human nasal secretions [73, 74].

### Chemical complementation of *dat*::Tn with D-alanine restores human nasal colonization to parental levels

Both *dat*::Tn and Δ*alr1* displayed colonization defects on HNO-ALI (**Fig. 3B**) and D-alanine is the common product of their encoded enzymes (**Fig. 1C**). Therefore, we hypothesized that *dat*-dependent generation of D-alanine is required for parental levels of nasal colonization. To test this, we exogenously supplemented the *dat*::Tn mutant at the time of inoculation onto HNO-ALI separately with each of the D-amino acids that Dat can produce: D-alanine and D-glutamate. To assess for possible dose-dependent effects during nasal colonization, we supplemented with a range of concentrations from 0.375 to 37.5 mM. As a control, we supplemented the parental *S. aureus* strain with the maximum dose of each D- and L-amino acid (37.5 mM) to assess whether this affected colonization levels independent of *dat*.

Colonization by the parental strain was unaffected by supplementation with either D or L -alanine or -glutamate (**Fig. 4**, left side), implying that *S. aureus* might be fine-tuned for life with the existing levels of L-glutamate and L-alanine on human nasal mucosa. Because there was no difference among the parental strain CFUs/HNO-ALI under the various conditions, we compared each *dat*::Tn condition to the unsupplemented (no added amino acids) parental condition. As in our previous experiments, the *dat*::Tn mutant alone (without amino acid supplementation) had a 7,208-fold colonization defect compared to the parental strain (**Fig. 4**: right side, gray).

**Figure 4.**
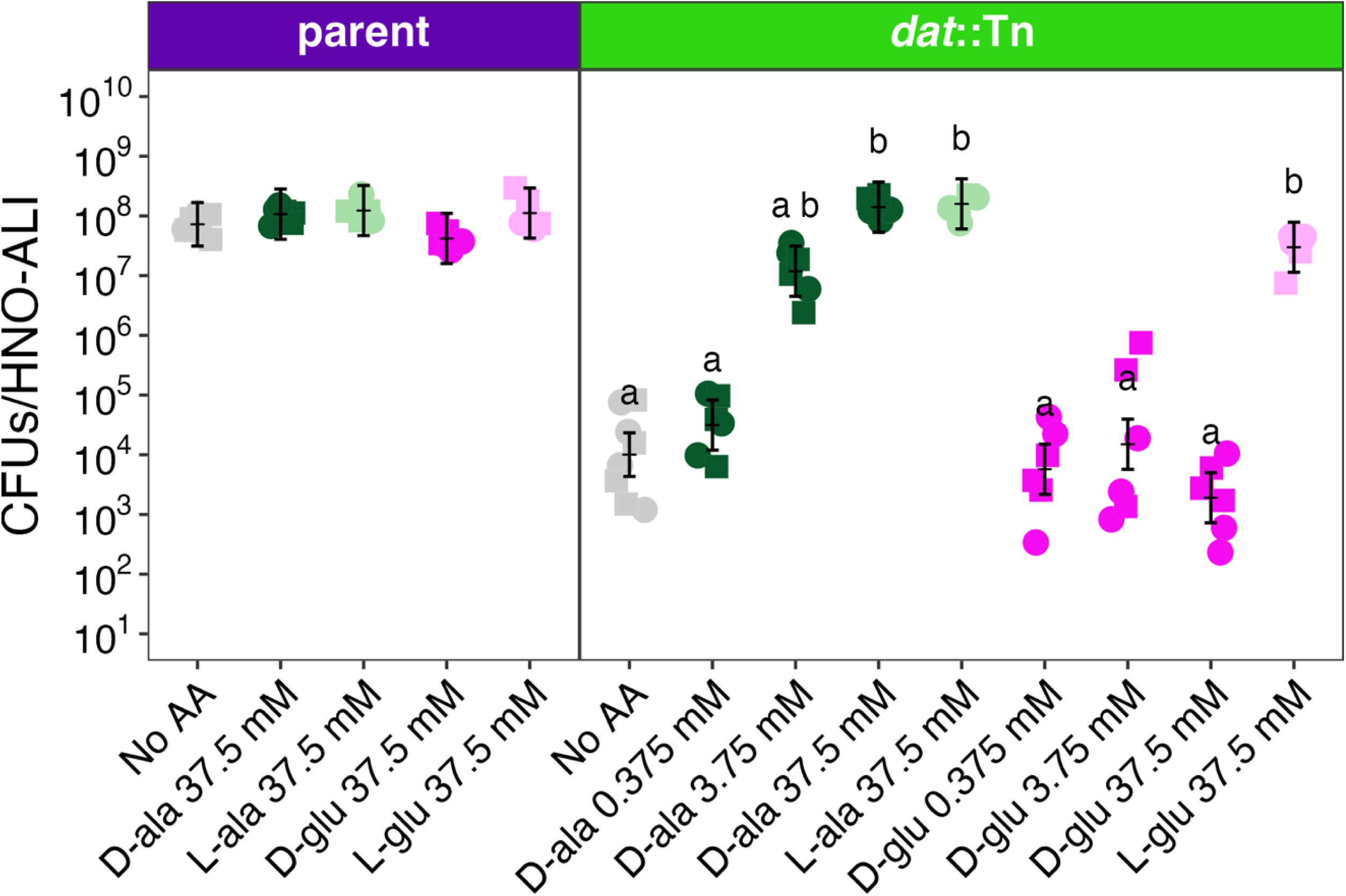
Exogenous D-alanine but not D-glutamate restores *S. aureus dat*::Tn colonization to parental levels on HNO-ALI. The parent or *dat*::Tn strains were colonized on HNO-ALI for 24 h and supplemented with either EBSS (gray) or D-alanine (dark green), L-alanine (light green), D-glutamate (dark pink), or L-glutamate (light pink) resuspended in EBSS in *n* = 3 independent experiments in each of 2 HNO lines (HNO204, circles and HNO918, squares). Log-transformed CFUs/HNO-ALI between groups were compared using an LMM with strain/media condition as a fixed effect and experimental date as a random effect. Pairwise contrasts were performed to compare all conditions to the unsupplemented parent strain (labeled “a” if Holm-adjusted *p* < 0.05) and to compare all *dat*::Tn conditions to the unsupplemented *dat*::Tn strain (labeled “b” if Holm-adjusted *p* < 0.05). Vertical brackets represent model-predicted mean values and confidence intervals (± 2 × SEM), back-transformed to the original response scale. Statistical analysis is in **Table S2C**.

Supplementation of *dat*::Tn with exogenous D-alanine resulted in a dose-dependent increase in colonization (**Fig. 4**: right side, dark green). Supplementation with 0.375 mM D-alanine had no effect. However, supplementation with 3.75 mM D-alanine was sufficient to increase the *dat*::Tn CFUs/HNO-ALI 1,180-fold and 37.5 mM D-alanine was sufficient to restore the *dat*::Tn mutant to parental-level colonization (13,986-fold higher than *dat*::Tn without amino acids). These data indicate that Dat plays a crucial role in the generation of D-alanine during human nasal colonization.

In contrast, supplementation with D-glutamate failed to rescue the *dat*::Tn mutant colonization defect on HNO-ALI, with no statistically significant difference in the CFUs/HNO-ALI of *dat*::Tn after 24 h of colonization without or with (0.375 up to 37.5 mM) exogenous D-glutamate (**Fig. 4**: right side, dark pink). This indicates that *dat-*dependent generation of D-glutamate is not limiting during colonization of a human nasal mucosal surface. MurI can also produce D-glutamate (**Fig. 1C**), which is required for peptidoglycan synthesis, since it is the second amino acid in the peptidoglycan stem peptide, and its position and chirality protect against cell wall degradation by host peptidases [75].

Unexpectedly, supplementing the *dat*::Tn mutant with 37.5 mM L-glutamate at the time of colonization restored it to parental levels of colonization (**Fig. 4**: right side, pale pink). We supplemented the *dat*::Tn mutant with L-glutamate to control for the possibility that *murI*-dependent interconversion of L- and D-glutamate might complement *dat*::Tn in the event that generation of D-glutamate was a primary role of Dat during colonization. However, as discussed above, our data indicate that the primary role for Dat during on colonization of HNO-ALI is generation of D-alanine. Therefore, we interpret complementation of the *dat*::Tn mutant colonization defect by exogenous L-glutamate (but not D-glutamate) as evidence that *S. aureus* can convert L-glutamate into L-alanine independent of *dat* (which could then be converted by Alr1 to D-alanine).

To test whether *alr1*-dependent production of D-alanine might complement the *dat*::Tn mutant for colonization, we also supplemented with exogenous L-alanine. Addition of 37.5 mM of L-alanine at the time of inoculation restored parental levels of colonization at 24 h (**Fig. 4**: right side, light green). This implies that *alr1* alone confers the capacity to produce sufficient D-alanine to support parental *S. aureus* colonization given a sufficient level of L-alanine as substrate. It also points to L-alanine as limiting in the human nasal mucosal habitat, in the absence of *dat*-dependent production of D-alanine.

### Disruption of *dat* reduces *S. aureus* nasal colonization fitness across strains from different clonal complexes

Our data show that *dat* is a colonization fitness factor in *S. aureus* LAC (a methicillin-resistant CC8 strain derived from a skin infection [46]) and its plasmid-free derivative JE2 [76]. We hypothesized that disruption of *dat* would also yield a greater colonization defect than disruption of *alr1* in another *S. aureus* strain. To test this, we transduced *dat*::Tn from our parent strain into two additional clinical isolates and a primary nasal isolate of *S. aureus*. We started with *S. aureus* 502A, a historical CC8 clinical isolate that was briefly used early in the 1960s for bacterial interference to therapeutically colonize infants and halt nursery outbreaks of invasive infection by a more virulent phage type 80/81 *S. aureus* strain [77]. We transduced *dat*::Tn or *alr1*::Tn in the 502A background and colonized HNO-ALI for 16 h with each strain alone and in competition with the parent. *S. aureus* 502A produces alpha-toxin but is described as less virulent than other CC8 *S. aureus* strains [77–79]. Therefore, we used the wild-type background (without adding an *hla*::Tn insertion). We colonized HNO-ALI with 502A for 16 h with minimal cytotoxicity (**Fig. S1C**) and recovered 107-to-108 CFUs/HNO-ALI of parental 502A in all conditions. In comparison, we recovered 17,410-fold fewer *dat*::Tn mutant CFUs and 177-fold fewer *alr1*::Tn mutant CFUs. (**Fig. S4**). Both the *dat*::Tn and *alr1*::Tn mutant were less fit than 502A, with competition indices significantly lower than 1 (**Fig. 5**). Similar to the *S. aureus* JE2 background, the *dat*::Tn CI of 0.00007 was 59-fold lower than that of the *alr1*::Tn CI of 0.0042 (*p* = 0.008), indicating that the predominant role of *dat* for fitness on HNO-ALI is not specific to a single strain.

**Figure 5.**
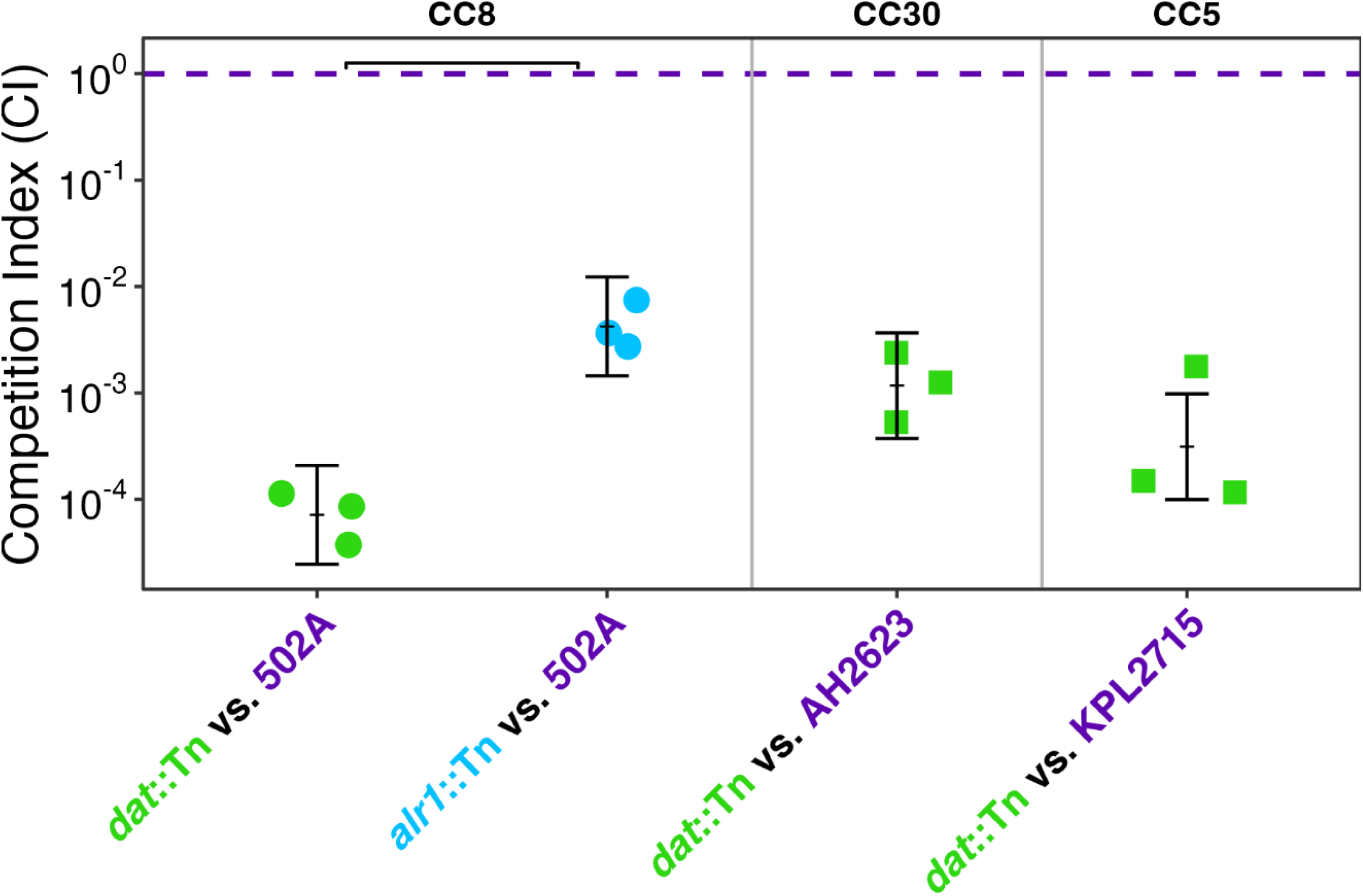
The *dat* gene is also more important than *alr1* for *S. aureus* fitness on human nasal mucosa in a second CC8 strain and *dat* contributes strongly to fitness in both CC5 and CC30 strains. Competitive indices (CI = (mutant output / parent output) / (mutant input / parent input)) of *dat*::Tn (green) or *alr1*::Tn (blue) in the indicated strain backgrounds: 502A (CC8), AH2623 (CC5), and KPL2715 (CC30) based on CFUs/HNO-ALI shown in **Fig S4**. *n* = 3 independent experiments in either of 2 HNO lines (HNO204, squares or HNO918, circles). Log-transformed CI were modeled with mutant as a fixed effect and experimental date as a random effect. A pairwise contrast was performed to compare mutants within the 502A strain background with a horizontal bracket representing statistically significant *p* < 0.05. Vertical brackets represent model-predicted mean values and confidence intervals (± 2 × SEM), back-transformed to the original response scale. The purple dashed line indicates a CI value of 1, representing equal fitness between the mutant and the parental strain, and serves as the reference value for the one-sample t-tests performed for each strain, all of which were statistically significant (*p* < 0.05). Statistical analysis is in **Table S2A**.

We further hypothesized that disruption of *dat* would substantially reduce fitness on HNO-ALI in strains belonging to other CCs. To test this, we transduced the *dat*::Tn insertion into *S. aureus* strains AH2623 [80] (a CC5 infection isolate) and KPL2715 (a CC30 primary human nasal isolate from our strain collection that is well tolerated by HNO-ALI at 24 h). We recovered 1,056-fold fewer CFUs *dat*::Tn compared to parent AH2623 after 16 h colonization, and 1,652-fold fewer *dat*::Tn CFUs compared to parent KPL2715 after 24 h colonization (**Fig. S4**). Colonization by either parent strain induced minimal cytotoxicity (**Fig S1C**). Again, in competition with the respective parent strain, the AH2623 *dat*::Tn mutant had a CI of 0.001, and the KPL2715 *dat*::Tn mutant had a CI of 0.0007 (**Fig. 5**) demonstrating a substantial loss in fitness in representative strains from two other CCs.

### The *S. aureus dat*::Tn mutant is unable to attain sufficient D-alanine from a cocolonizing nasal microbiont to rescue its colonization defect

D-alanine is an essential amino acid for bacteria in a mammalian host environment [81, 82]. It is produced internally then translocated across the cell membrane for incorporation into peptidoglycan and, in gram-positive bacteria, for decoration of teichoic acids on the cell surface. Similar to our data above with only *S. aureus*, most pathogenesis studies showing a requirement for D-alanine are done during monoinfection [41]. This raises the question of whether D-alanine might be a public good that can be scavenged from other bacteria in the microbiota. The 1,502-fold colonization defect of *dat*::Tn in competition with the parental *S. aureus* (**Fig. 2B**), showed that cocolonization with another *S. aureus* strain that is wild-type for D-alanine production is insufficient to exogenously complement the colonization defect. Based on this, we hypothesized that another microbiont would also be an insufficient source of exogenous D-alanine in the nasal mucosal environment. To test this, we cocolonized HNO-ALI with *S. aureus dat*::Tn (in the JE2 *hla*::Tn background) and *Corynebacterium pseudodiphtheriticum* KPL1989 [83]. *C. pseudodiphtheriticum* is part of the nasal microbiota in about 20% of adults [84]. Cocolonization was insufficient to rescue the colonization defect of an *S. aureus dat*::Tn mutant, and it did not impact parental *S. aureus* colonization (**Fig. 6**), further supporting that there is insufficient D-alanine from another common bacterium in the nasal environment to compensate for loss of *dat* function.

**Figure 6.**
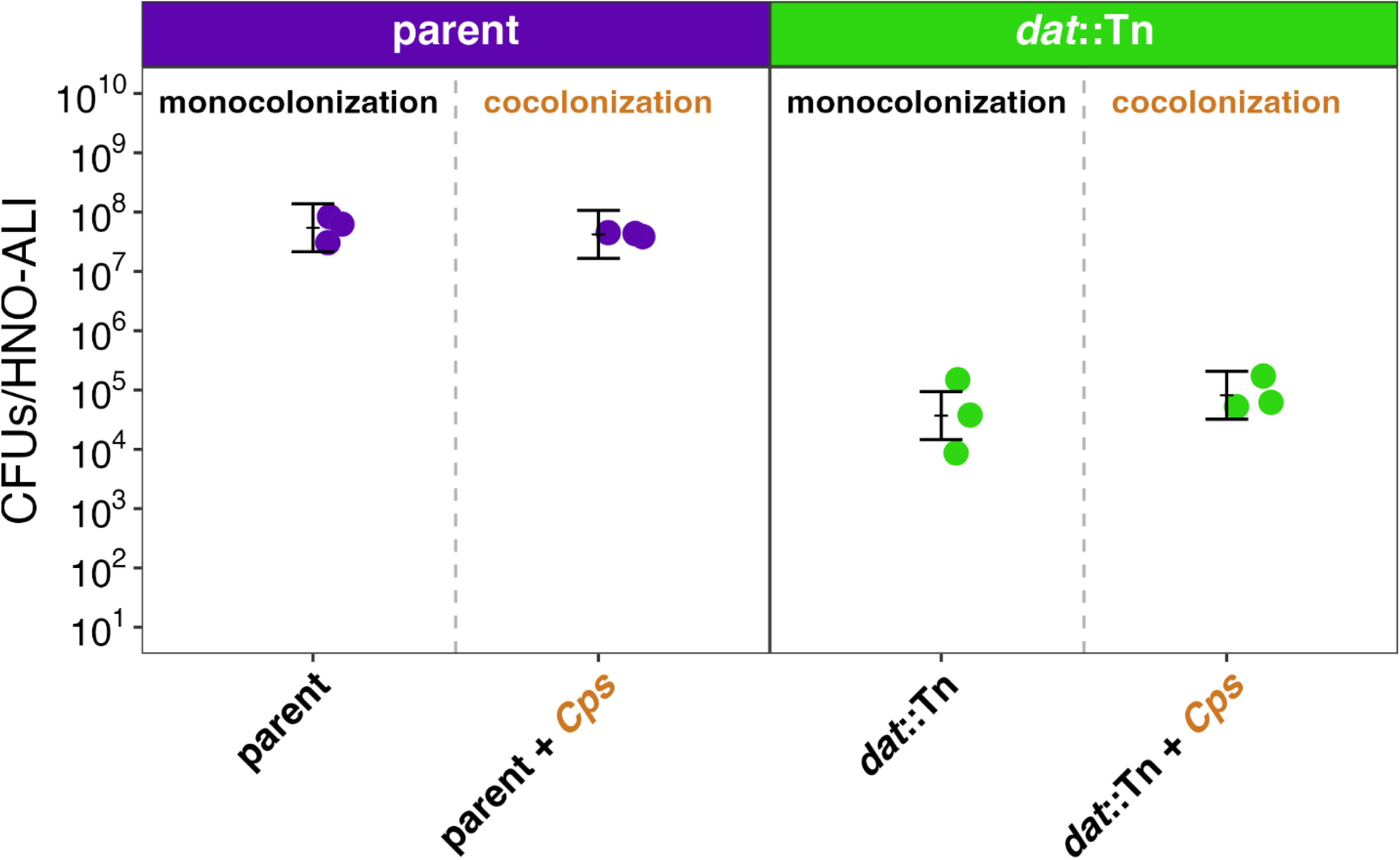
Cocolonization with *Corynebacterium pseudodiphtheriticum* is insufficient to rescue an *S. aureus dat*::Tn mutant. HNO-ALI were cocolonized with 10^6^ CFUs/HNO-ALI of *C. pseudodiphtheriticum* plus 10^5^ CFUs/HNO-ALI of either the parent *S. aureus* KPL4530 (purple) or the *dat*::Tn mutant (green) for 24 h. *n* = 3 independent experiments in HNO918. Log-transformed CFUs/HNO-ALI were compared between groups using an LMM with strain and colonization condition as fixed effects and experimental date as a random effect. Pairwise contrasts were performed to compare colonization conditions within each strain, and no significant differences were observed. Additional contrasts comparing *dat*::Tn vs. parent were significant under both monocolonized and uncolonized conditions (*p* < 0.05). Vertical brackets represent model-predicted mean values and confidence intervals (± 2 × SEM), back-transformed to the original response scale. Statistical analysis is in **Table S2D.**

## DISCUSSION

Together, the data presented in this study indicate that *dat-*dependent production of D-alanine is crucial for fitness during colonization of a human nasal mucosal surface. Cumulatively, these results are consistent with the human nasal mucosal environment imposing a strong positive selective pressure for the maintenance of a functional *dat* gene in *S. aureus*.

The predominance of *dat* over *alr1 ex vivo* on HNO-ALI shown here contrasts with the established predominance of *alr1* over *dat* for production of D-alanine *in vitro* in culture medium [36]. Based on *in vitro* data, prior studies point to Alr1 as a potential antimicrobial therapeutic target against *S. aureus* [37] and other bacterial species [38–40]. The findings detailed here suggest targeting *dat*, for example with antimicrobials such as β-Chloro-D-alanine (BCDA) [85], would be a more favorable strategy for inhibiting *S. aureus* during human nasal decolonization.

The composition of amino acids and oligopeptides available in human nasal mucus may influence the relative importance of *dat* and *alr1* during *S. aureus* nasal colonization. Specifically, *S. aureus’s* reliance on Dat more than Alr1 to generate D-alanine suggests that L-glutamate is more abundant while L-alanine is limiting. Two studies describe the metabolomic composition of *in vivo* human nasal secretions. Krismer et al. reports similar concentrations of free alanine (19.0-to-248.0 µM) and free glutamate (39.8-to-103.4 µM) [73] in human nasal secretions from eight donors. Detection of oligopeptides was not reported. However, *S. aureus* also uses oligopeptides as a source of sulfur [86] and amino acids [87]. Farne et al. also reports comparable levels of free alanine and glutamate in metabolomic profiling of human nasal secretions from eight donors [74]. In addition, they detected a variety of oligopeptides in nasal secretions, many of which contained γ-glutamic acid. For example, γ-glutamyl di- and tri-peptides, including glutathione (γ-glutamyl-L-cysteine-L-glycine), each of which were present in a relative concentration similar to that of free glutamate (based on their supplemental data) [74]. Elsewhere, glutathione is described as abundant in human olfactory mucus [88]. These data point to the possibility that *S. aureus* might use oligopeptides rich in glutamate as a key amino acid source in human nasal mucus. If so, this may allow *S. aureus* to rely more on Dat than on Alr1 for production of D-alanine, consistent with the more severe colonization defect we observed in a *dat*::Tn mutant. Our screen also identified some oligopeptide importers and peptidases as candidate fitness genes on HNO-ALI, e.g., highlighted in blue in **Table S1A**. Since *S. aureus* imports oligopeptides [87], including glutathione [88], the potential fitness role of peptidases during human nasal mucosal colonization points to an importance of *S. aureus* breaking down oligopeptides in this environment. While we propose that the amino acid composition of human nasal secretions drive the colonization defect seen in a *dat*::Tn mutant, additional studies are needed to support this possibility.

This study provides evidence for a possible promoter 5’ of *dat* that overlaps with the 3’ coding sequence of the preceding *pepV* gene in the bicistronic operon. The P*datdat* construct (which includes 317 bp 5’ of *dat*) restored the *dat*::Tn colonization defect to near-parental levels *ex vivo* on HNO-ALI. In contrast, the P*datdat* construct failed to rescue the *dat*::Tn growth defect *in vitro* in defined medium lacking alanine (CDM-ala). These data point to differential regulation of the putative P*dat* in response to environmental cues that vary between HNO-ALI and this medium, for example, availability of glucose or oligopeptides and amino acids. *S. aureus* amino acid synthesis is controlled by catabolite repression [89] and glucose in human nasal secretions is limiting [73, 90]. This is an area for future investigation, since a defined P*dat* would permit future studies on its regulation in response to specific environmental cues.

D-alanine synthesis via *alr1* and/or *dat* is important for *S. aureus* resistance to antibiotics and for fitness during infection. Loss of D-alanine synthesis in defined media (*alr1*) and import (*cycA*, the primary alanine importer) increases susceptibility to β-lactam antibiotics, D-cycloserine, and a variety of other antimicrobial compounds [37]. A *dat alr1* double mutant is auxotrophic for D-alanine, and a *dat alr1 alr2* triple mutant has reduced virulence in murine models of bacteremia and subcutaneous-catheter, device-associated infection and has been proposed for use in vaccines [41]. Additionally, *S. aureus* lacking *dat* and *murI* (a D-glutamate auxotroph) is cleared in a murine bacteremia model without causing disease [91]. These data suggest that maintaining D-alanine pools is advantageous to *S. aureus* during antibiotic treatment of infection, as it is known to be important for resistance to organic acid intoxication [36].

To our knowledge, this is the first study using an *ex vivo* model of human nasal respiratory epithelium and TnSeq to identify candidate *S. aureus* fitness factors for nasal colonization. Lyon et al. report *alr1* as a candidate fitness gene on mouse cervix (but not other parts of the vaginal tract) [24]. In contrast, *dat* was not detected as a fitness gene during mucosal colonization of the mouse vaginal tract. This might reflect differences in human vs. murine mucosal environments and/or in vaginal vs. nasal mucosal environments, and emphasizes roles for both *in vivo* animal models, which can query effects of an intact immune system, and *ex vivo* human epithelial models, which query human-specific environmental factors, to study bacterial colonization and microbiota.

Apart from *alr1*, many candidate fitness factors identified on nasal mucosa overlap with factors identified during murine vaginal colonization, many of which are genes related to metabolism [24]. For example, *cntE*, *ccpA*, *purE, codY, aroAB, and menBCDE* contribute to *S. aureus* fitness during mucosal colonization of both the murine vaginal and human respiratory tract (**Table S1A**). Additionally, *cntE*, *purE*, *aroBD*, and *menBD* were important for *S. aureus* intracellular infection of A549 cells [19].

Purine biosynthesis pathways in particular were significantly enriched for genes contributing to fitness in our study (**Fig. 1B**) and during murine vaginal colonization [24]. The importance of purine biosynthesis and scavenging is well established as a core metabolomic process in the context of infection [20, 59–62]. Here, we found that both *de novo* purine biosynthesis (*purB, purD*, *purE,* and *purH*) and uptake (*apt, xpt, hpt*) promote colonization fitness on HNO-ALI. The overlap between identified fitness factors in an *in vivo* mouse model of vaginal mucosal colonization and a reductionist *ex vivo* model of human nasal mucosal colonization (HNO-ALI) supports the broader importance of these genes for *S. aureus* fitness during mucosal colonization.

Finally, it is interesting that exogenous addition of L-glutamate complemented the *dat*::Tn colonization defect, but D-glutamate did not. Since *S. aureus* imports D-glutamate (a D-glutamate auxotrophic strain of *S. aureus* is complemented with exogenous D-glutamate [91]), this finding points to Dat generation of D-alanine, rather than D-glutamate, as the most crucial during nasal colonization. Since Alr1 conversion of L-alanine to D-alanine can supply D-alanine in a *dat*::Tn mutant, the L-glutamate rescue points to a metabolic pathway by which *S. aureus* converts L-glutamate to L-alanine. In other bacterial species these are interconverted by L-alanine aminotransferase. For example, *Bacillus subtilis* encodes alanine transaminase *alaT* [92]. However, to our knowledge, *S. aureus* lacks a specific L-alanine aminotransferase (and did not encode a protein with high homology to *B. subtilis* AlaT when we searched via BLASTp). Aminotransferases are well described to have substrate flexibility [93], so it is possible that one of the known *S. aureus* aminotransferases can serve this role in the absence of *dat*. Additionally, there are more circuitous biosynthetic routes from L-glutamate to L-alanine. For example, *gudB* converts L-alanine to 2-oxoglutarate, which feeds into the tricarboxylic acid cycle and generates pyruvate. If pyruvate is abundant, alanine dehydrogenases (Ald1 and Ald2) can then generate L-alanine. Finally, the *dat*::Tn rescue by L-glutamate but not D-glutamate also implies that MurI did not convert a sufficient amount of imported D-glutamate to endogenous L-glutamate to rescue to the *dat*::Tn colonization defect on HNO-ALI.

Use of an early colonization timepoint (8 h) to identify *S. aureus* colonization fitness factors is a limitation, since there may be other fitness factors important for sustained nasal colonization. In addition, although HNO-ALI are an excellent model system for human nasal respiratory epithelium, it is a reductionist system without immune cells or the full complexity of the human nasal microbiota.

Overall, this study strongly supports *dat*-dependent production of D-alanine as crucial for *S. aureus* fitness on the human nasal mucosa that coats the majority of the nasal passages, and where *S. aureus* is known to colonize [27]. Thus, Dat may be worth considering as a possible target to improve nasal decolonization. It also identified a putative promoter, the study of which might provide new insights into *S. aureus’* response to the human nasal mucosal environment early in colonization. More broadly, this study demonstrates a use for HNO-ALI to identify human-specific colonization fitness factors in bacteria.

## MATERIALS and METHODS

### HNO-ALI culture

Adult-derived HNOs from three different donors were propagated as 3D organoids (as we previously described [31]) and were similarly differentiated at an ALI, except that the BCM 3D Organoid Core made a few modifications to their protocol to improve HNO-ALI consistency and reliability [32]. These modifications include the following: 1) during 3D HNO culture, the Airway Organoid Propagation medium now includes 30% Wnt-conditioned medium (made by growing the ATCC cell line L Wnt-3A (#CRL-2647) in Advanced DMEM/F12 (ThermoScientific #12634-028) + 10% FBS + 1x Glutamax (ThermoScientific #35050061)), and 2) monolayers for HNO-ALI are now seeded using 1.5 x 105 cells per Transwell® insert (Corning #3470). Apical airway organoid (AO) medium was removed after 72 h. During HNO-ALI colonization, 600 µL of PneumaCult Airway Organoid Differentiation Medium (AODM; STEMCELL Technologies #05060) is used on the basal side of the transwell inserts.

One day before bacterial colonization (day -1) we replaced the 600 µL of spent basal AODM in each well with fresh AODM, performed quality control measurements as detailed below, and put HNO-ALI at 34 °C in a humidified, 5% CO2-enriched incubator to equilibrate to nasal passage temperature [47] for 24 h before colonization. Basal medium was again replaced at -1 h before colonization.

### HNO-ALI quality control

In addition to the standard quality control measures routinely performed by the BCM 3D Organoid Core, we performed the following on one transwell per experiment to ensure the quality of each HNO-ALI experiment (as previously described [31]): 1) measured transepithelial electrical resistance (TEER), with a minimum (range) TEER value across all included experiments of 700 (range 700 - 1250) Ohms; 2) counted the number of epithelial cells per HNO-ALI, with a range of 172,600 – 902,000 cells; 3) tested for contamination with a PCR-based *Mycoplasma* detection assay. Briefly, after measuring TEER, HNO-ALI cells were scraped from the transwell with a pipette tip and resuspended in the 100 µL of Earle’s Balanced Salt Solution without calcium, magnesium, or phenol red (Gibco™ #14155063) (EBSS) used to measure TEER. Cells were then heated to 95 °C for 10 min to lyse eukaryotic cells. The lysate was used for PCR-based *Mycoplasma* detection as per the manufacturer’s instructions (Biovision Mycoplasma PCR Detection kit Abcam #AB289834) followed by electrophoresis in a 1% agarose Tris Acetate EDTA (TAE) gel (Rockland Immunochemicals #MB-020) and visualization with SYBR Safe (Invitrogen #S33102).

### Ethics statement

For experiments in this study, we purchased de-identified, anonymized HNO-ALI from the BCM 3D Organoid Core on a fee-for-service basis. The Baylor College of Medicine (BCM) 3D Organoid Core has established HNOs with approval from the BCM Institutional Review Board and with donor informed consent under protocol H-46014 [31].

In a prior study, we isolated bacterial strains via swabbing the nasal vestibule (nostrils) of generally healthy children and adults in the United States in 2017, 2018, and 2019 under protocol #17-02 approved by the Forsyth Institute Institutional Review Board [72, 94]. All adults provided written informed consent and all children 5 years and older provided assent in addition to the written informed consent provided by their parent and/or guardian.

### Primary nasal bacterial isolation

*S. aureus* strain KPL2715 (passaged as KPL4360, see **Table S4**) was isolated from a human nostril swab on BHI supplemented with 1% Tween80 and 25 µg/mL fosfomycin according to our established isolation protocols with written parental consent from an assenting child aged 7 to 12 years [72, 94]. Whole genome sequencing and taxonomical assignment was performed by the former Microbial Genome Sequencing Center and deposited in Bioproject PRJNA842433.

### Bacterial strains and *in vitro* and *ex vivo* culture conditions

**Table S4** lists all bacterial strains and plasmids used. The Nebraska Transposon Mutant Library (NTML) NR-48501 [76] and the NTML Toolkit [95] were shared with us via the Biodefense and Emerging Infections Research Resources Repository (BEI Resources) (www.beiresources.org) from the Network on Antimicrobial Resistance in *S. aureus* Repository (NARSA). For HNO-ALI inoculation, each bacterial strain of interest was struck for single colonies on a sterile 47-mm-diameter, 0.2-μm polycarbonate membrane (Millipore Sigma #GTTP04700) atop tryptic soy 1.5% agar (TSA) medium and grown at 34 °C overnight (16 - 20 h) in a humidified, 5% CO2-enriched incubator. We then picked 3-4 single colonies from each membrane using a sterile p1000 pipette tip for resuspension in 1 mL EBSS. The optical density at 600 nm (OD600) was normalized to an OD600 equal to 105 CFUs in a 15 µL inoculum for monoculture conditions and to 105 CFUs in a 7.5 µL in cocolonization (competition) conditions.

For HNO-ALI colonization, 15 µL of bacteria were inoculated into the middle of each transwell insert without touching or disturbing the epithelium. The 24-well plates were tapped gently on the bench to distribute inoculum before gentle centrifugation at 120 rpm for 1 min in a RT Eppendorf 5430 R centrifuge equipped with an A-2-MTP windshield rotor to help distribute bacteria into the HNO-ALI mucus layer. Colonized HNO-ALI were incubated in a 34 °C humidified 5% CO2-enriched incubator for 24 h (unless otherwise stated) before collection. Bacterial CFUs were enumerated as described previously [31] and as below in the TnSeq screen experiment section.

### Cytotoxicity measured by Lactate Dehydrogenase (LDH)

For all HNO-ALI colonization experiments, basal medium was collected -1 h before colonization and after each experimental endpoint. Cytotoxicity was measured by quantifying lactate dehydrogenase (LDH) using a CytoTox Non-Radioactive Cytotoxicity Assay (Promega #G1780) per the manufacturer’s instructions in duplicate as described previously [31].

### TnSeq screen experiment

We used the *S. aureus* LAC TnSeq library generated by Melinda Grosser and colleagues [17], receiving an aliquot as a gift from Dr. Anthony Richardson. To generate new working stocks of the Tn library, we outgrew 1 mL of the library in 100 mL of fresh brain heart infusion (BHI) (Difco #237200) supplemented with 5 µg/ml erythromycin (erm) (Sigma-Aldrich #E5389-5G) shaking at 250 rotations per min (rpm) at 34 °C for 4 h. The resulting batch culture was aliquoted as 1 mL portions in microcentrifuge tubes and centrifuged at 10,000 x *g*. Each pellet was resuspended in BHI with 20% glycerol (Thermo Fisher #J61059.K2), flash-frozen in an isopropanol and dry ice bath, and stored at −80 °C. For HNO-ALI inoculation, we thawed an aliquot on ice, pelleted it by centrifugation at 10,000 x *g* for 10 min, and resuspended in EBSS, adjusting to a final OD600 equal to 10^6^ CFU in a 15 µL inoculum. On two separate dates, we colonized HNO-ALI, line HNO919, with the Tn library and separately with *S. aureus* LAC as a control.

CFUs were collected from HNOs after 8 h colonization in a humidified, 5% CO2-enriched incubator at 34 °C, as described previously [31]. Briefly, each complete HNO-ALI transwell was removed from the basal medium, treated with 75 µL 0.25% Trypsin-EDTA (Gibco™ #15400054), and incubated at 34 °C for 15 min. Then, 75 µL of 0.025% Triton X-100 (Thermo Scientific™ #J66624.AE) was added to each transwell and eukaryotic cells were lysed mechanically by vigorously pipetting up and down 30 times. Each sample was then serially diluted in EBSS and 5 µL of each dilution was spotted on BD BBL™ Columbia agar medium with 5% sheep’s blood (Becton, Dickinson and Company #90006-198-CS) to enumerate CFUs.

For TnSeq, each dedicated HNO-ALI sample was harvested as above. The complete trypsin- and Triton X-100-treated HNO-ALI sample was subsequently inoculated into 5 mL of fresh BHI and cultured for 3 - 4 h shaking at 250 rpm at 34 °C to allow for bacterial outgrowth. Cultures were then centrifuged at 10,000 x *g* for 10 min at 4 °C, the supernatants were discarded, and the resulting cell pellets were flash-frozen in an isopropanol and dry ice bath and stored at −80 °C until further processing.

### DNA extraction and TnSeq library preparation for sequencing

Bacterial cell pellets were thawed on ice and DNA was extracted using the Biosearch Technologies MasterPure Gram Positive DNA Purification Kit (#MGP04100) with the following modifications. Prior to extraction, cell pellets were washed by resuspension in 5 mL TES buffer (tris(hydroxymethyl)methyl-2-aminoethanesulfonic acid) and centrifuged at 5,000 x *g* for 10 min at 4 °C. The resulting pellets were resuspended in 300 µL TE buffer (kit). Lysostaphin (2 µL of a 10 µg/µL freshly prepared stock) (Sigma Aldrich #L7386) was added to each sample and incubated at 37 °C for 20 min to facilitate cell wall digestion. Then, 300 µL of Gram-Positive Lysis Solution (kit) was added. After vortexing for 10 sec, samples were transferred to Lysing Matrix B tubes (M.P. Biomedicals #1169110-CF) for mechanical disruption using a FastPrep-24™ 5G (M.P. Biomedicals #11-600-5500) at 6.0 m/s for four cycles of 30 s each, with 30-second rest intervals on ice between cycles.

Following bead beating, samples were treated with 2 µL Proteinase K (kit) at 65 °C for 15 min, then equilibrated to room temperature (RT) for 5 min. Protein precipitation was performed by adding 350 µL MPC Protein Precipitation Reagent (kit) to each sample, vortexing for 10 sec, and centrifuging at >10,000 x *g* for 10 min at 4 °C. The clarified aqueous layer was transferred to a clean 1.5 mL microcentrifuge tube and treated with 2 µL RNase A (kit) for 30 min at 37 °C to remove RNA.

DNA was precipitated by addition of 700 µL sterile isopropanol followed by gentle mixing with 30 - 40 inversions. Precipitated DNA was pelleted by centrifugation at >10,000 x *g* for 10 min at 4 °C. The isopropanol was removed, and the DNA pellet was washed twice with 1 mL 70% ethanol, air-dried for 15 min at RT, and resuspended in 100 µL nuclease-free water. DNA concentration and quality were assessed with a Qubit™ 4 Fluorometer(Invitrogen™ #Q33226) using the AccuGreen™ Broad Range dsDNA Quantitation Solution (Biotium #31069T). Genomic DNA samples were stored at 4 °C until downstream processing.

Purified genomic DNA was normalized to a concentration of 50 ng/µL and sheared to a target fragment size of ∼350 bp [22, 50] by sonication using a Covaris LE220 Focused Ultrasonicator with the following parameters: peak power, 450 W; duty factor, 15%; cycles per burst, 200; total treatment time, 63 s. Sheared samples were imaged on a 1.5% agarose TAE gel using SYBR™ Safe DNA Gel Stain (Invitrogen, #S33102) to ensure DNA fragments were between approximately 200 and 500 bp.

To facilitate selective amplification of Tn-containing fragments, sheared DNA was subjected to terminal deoxynucleotidyl transferase (TdT) poly-C tailing (New England BioLabs #M0315L). Each reaction contained 50 µL sheared DNA (2.5–5 µg), 5 µL of a ddCTP/dCTP mixture (0.5 mM ddCTP / 9.5 mM dCTP), 1 µL TdT enzyme, 7.5 µL 10X TdT buffer, 7.5 µL CoCl2, and 4 µL nuclease-free water, for a total reaction volume of 75 µL. Reactions were incubated at 37 °C for 90 min with shaking, and mixed by gentle flicking every 30 min. Tailed DNA was purified using the QIAquick PCR Purification Kit (Qiagen #28104) and quantified on a Qubit™ 4 Fluorometer with the AccuGreen™ Broad Range dsDNA Quantitation Assay.

Selective enrichment of Tn-containing fragments was achieved through a two-round PCR strategy [50] using the method of Ibberson et al. [22]. Adapted primer sequences are listed in **Table S5**. In the first PCR (PCR1), a biotinylated primer specific to the mariner Tn sequence, oKL922 (a modified version of SA-PCR1-BA [22]), was paired with a non-specific primer complementary to the poly-C tail, oKL923 (a modified version of olj376 [96]), to enrich for and biotinylate DNA fragments containing a Tn insertion. We used the Expand Long Template DNA polymerase (Roche, cat. no. 11681842001) for PCR reaction enzymes. Each 50 µL reaction contained 200 ng TdT-tailed DNA, 5 µL 10X Roche Buffer 2, 2.5 µL 10 mM dNTPs, 2 µL 30 µM oKL922, 3 µL 30 µM oKL923, and 0.8 µL Roche DNA polymerase, with nuclease-free water to volume. Up to 16 reactions were performed per sample. Thermocycler conditions were as follows: initial denaturation at 95 °C for 5 min; 10 cycles of 94 °C for 30 s, 55 °C for 30 s, and 68 °C for 2 min; final extension at 68 °C for 10 min. PCR1 products from each sample were pooled and purified using the Qiagen PCR cleanup kit. If samples were too alkaline following purification, sodium acetate was added to neutralize pH prior to cleanup, per kit instructions. Purified PCR1 products were quantified using a Qubit dsDNA Broad Range Assay. Biotinylated PCR1 products were captured on streptavidin-coated magnetic beads (Dynabeads M-270; Thermo Fisher Scientific) to remove non-specific products, per the manufacturer’s instructions. Beads were equilibrated to RT prior to use. For each reaction, 32 µL of beads were washed three times with 1 mL 1X Binding and Washing (B&W) buffer (2X stock: 2 M NaCl, 10 mM Tris-HCl, 1 mM EDTA, pH 7.5) on a magnetic stand, then resuspended in 50 µL 2X B&W buffer. An equal volume (50 µL) of purified PCR1 product (125–200 ng) was added to the resuspended beads, and samples were incubated at RT for 30 min with rotation on a tabletop mini orbital shaker (VWR #12620-938) at 250 rpm. Bead-bound DNA was washed once with 100 µL 1X B&W buffer and twice with 100 µL LoTE buffer (3 mM Tris-HCl, 0.2 mM EDTA, pH 7.5), then retained on the magnetic stand with the final wash removed.

A second PCR (PCR2) was performed directly on the bead-bound template to amplify DNA sequences and to introduce Illumina P5 and P7 sequencing adaptors. Each 50 µL reaction was assembled directly on the bead pellet and contained 5 µL 10X Roche Buffer 2, 3 µL 10 mM dNTPs, 2 µL 30 µM Tn-specific primer (oKL924; SA-Mariner-PCR2 [22]), 2 µL 30 µM reverse primer (oKL925), and 0.8 µL Roche DNA polymerase, with nuclease-free water to volume. To preserve both DNA and enzyme integrity, the polymerase was added after bead resuspension and reactions were mixed by gentle tapping only. Vortexing was avoided to prevent shearing of bead-bound DNA. A minimum of four PCR2 reactions were performed per sample and pooled for later steps. Thermocycler conditions were: 95 °C for 5 min; 15 cycles of 94 °C for 30 s, 58 °C for 30 s, and 68 °C for 2 min; final extension at 68 °C for 10 min. Reactions were gently mixed twice during cycling, with the thermocycler paused during the 68 °C extension step to facilitate gentle mixing. Following amplification, tubes were placed on a magnetic stand to separate the beads and the supernatant, containing the PCR2 product, was transferred to a new tube. PCR2 products were purified using a Qiagen PCR cleanup column to concentrate the sample and normalize pH prior to further cleanup.

Final DNA library size selection was performed using AMPure XP beads (Beckman Coulter #A63880) to remove residual primers, adapter dimers, and small fragments. AMPure beads were equilibrated to RT and vortexed for 30 s immediately prior to use. Sample volume was adjusted to 50 µL with nuclease-free water, and beads were added at a 0.9X volumetric ratio (45 µL) for the first cleanup round. Samples were mixed thoroughly by pipetting and incubated at RT for 15 min, shaking at 250 rpm on a tabletop mini orbital shaker (VWR #12620-938). Tubes were placed on a magnetic stand for 3 min to clear the suspension, the supernatant was discarded, and the bead pellet was washed twice with 800 µL freshly prepared 80% ethanol, with 30 s incubations per wash. Residual ethanol was removed, and the pellet was air-dried on the magnetic stand for 3-5 min, until the pellet transitioned from a glossy to a matte appearance. The pellet was resuspended in nuclease-free water and incubated at RT with shaking for 5 min, after which the eluate was recovered by magnetic separation and transferred to a new tube. This AMPure cleanup was performed twice in succession to ensure removal of adapter contamination. Final library concentration and quality was determined using an Agilent TapeStation (Agilent Technologies #4150) and TapeStation High Sensitivity DNA kit reagents (Agilent Technologies #5067) to ensure there were no DNA fragments smaller than ∼100 bp prior to sequencing. Samples were stored at 4 °C until submission for sequencing.

### Tn Sequencing

TnSeq libraries were sequenced as a fee-for-service by SeqCoast Genomics. Libraries were generated using the Illumina DNA Prep kit (Illumina #20060059) with Unique Dual Indexes omitting tagmentation and modifying bead cleanup to retain longer fragments. Sequencing was performed on an Illumina NextSeq2000 platform (2×150 bp), 10 million reads per sample, with 2% PhiX spike-in and on-instrument demultiplexing and trimming using DRAGEN v4.2.7.

### TnSeq read processing and insertion site mapping

TnSeq data were analyzed using an adapted version (https://klemonlab.github.io/DatCol_Manuscript/Manuscript_DatCol_Tn-Seq_Pipeline.html) of a previously published pipeline [22, 52]. Reads were mapped to a curated *Staphylococcus aureus* USA300_FPR3757 reference genome containing only the chromosomal sequence (NC_007793.1). All TA sites were identified computationally to generate a TA-site reference set.

Tn-specific inverted repeat sequences were removed using cutadapt [97] , and trimmed reads were aligned to the reference genome with bowtie2 [98, 99]. Unmapped and supplementary alignments were removed with samtools [100]. TA insertion sites were then quantified to generate per-sample tables of insertion coordinates and read counts.

### Identification of underrepresented Tn insertion sites

Identification of underrepresented Tn insertion sites in genes followed a Monte-Carlo simulation-based framework adapted from prior work [22, 52]. For each condition, insertion-site count tables were merged across replicates and normalized using DESeq2 size-factor estimation [101]. To model the expected distribution of non-essential insertions, 100 simulated datasets were generated by sampling from all possible genomic TA sites and from the empirical distribution of observed read depths.

Observed and simulated (expected based on all possible TA sites) insertion counts were binned into genes using the TnGeneBin.pl script, producing per-gene read counts and the number of insertion sites. Differential fitness was assessed by comparing observed vs. simulated (expected) counts using DESeq2, generating log2 fold-change (log2FC) estimates and adjusted *p*-values.

To classify essential genes, log2FC distributions were modeled using a two-component Gaussian mixture model (mclust) [102]. Genes assigned to the reduced-fitness category were designated candidate fitness factors if they had low classification uncertainty (≤ 0.01 or ≤ 0.05) and significant adjusted p-values (≤ 0.01 or ≤ 0.05).

### Generation of a markerless deletion of *S. aureus alr1*

*E. coli* strains were grown on Luria-Bertani, Miller, (LB) agar (BD Difco™ 244520) or shaking at 250 rpm in LB broth (BD Difco™ 244620) at 37 °C, with 100 µg/mL ampicillin (amp) (Ward’s Science #470233-552) as needed for plasmid selection and maintenance.

*S. aureus* strains were grown on TSA or shaking at 250 rpm in TSB containing 10 µg/mL chloramphenicol (cam) (Sigma-Aldrich #C0378).

Plasmid pKQ002 was constructed by Gibson Assembly into the SmaI site of the temperature sensitive allelic exchange vector pJB38 [95]. A ∼650 bp region 5’ of *alr1* (SAUSA300_RS11155) was PCR-amplified using Phusion™ High-Fidelity Master Mix (NEB M0530S) with primers oKL940 and oKL941. An ∼800 bp 3’ of *alr1* was amplified with primers oKL942 and oKL943. The pJB38 backbone was linearized with SmaI (NEB R0141S) following the manufacturer’s instructions. Gibson Assembly (NEB E5510S) was performed using a 5-fold molar excess of each fragment relative to the linearized pJB38. Assembly reactions were transformed into chemically competent *Escherichia coli* DH5α (NEB 5-α), per the manufacturer’s instructions. The resulting plasmid, pKQ002, contains the 5’ and 3’ homology regions required to generate a markerless deletion of *alr1* via allelic exchange in *S. aureus*. *E. coli* transformants carrying pKQ002 were colony purified three times on LB amp agar prior to preparation of frozen stocks.

Allelic exchange was performed as described in [95]. For plasmid propagation, a single DH5*α* colony carrying the plasmid of interest (KPL4565) was inoculated into 5 mL LB amp, grown overnight, diluted 1:100 into fresh medium, and cultured to mid-exponential phase (2-3 h). Plasmid DNA was isolated using the QIAprep Spin Miniprep Kit (Qiagen #27104) and eluted in nuclease-free water.

We then electroporated plasmid pKQ002 (10-100 ng) into the restriction negative *E. coli* strain DC10B, as described below. DC10B colonies carrying pKQ002 were selected for and then colony purified thrice on LB amp agar. For transformation into *S. aureus* JE2, plasmid DNA was concentrated to 500 ng/µL using a Savant™ SpeedVac™ DNA 130 Integrated Vacuum Concentrator System (Thermo Scientific™ #DNA130) for 30 min at RT, then deionized by placing 20 µL of the plasmid atop a 0.025 µm MCE membrane (MF-Millipore™ #GSWP02500) floating on 35 mL sterile Milli-Q water for 15-30 min. Then, 5 µg of deionized plasmid DNA was electroporated into *S. aureus* JE2, as previously described [103]. Transformants were selected on TSA cam, colony-purified thrice, stocked in LB + 15% glycerol, and frozen at −80 °C. Carriage of pKQ002 was confirmed by PCR using primers oKL926 and oKL927.

For allelic exchange, transformed *S. aureus* JE2 carrying pKQ002 (KPL4532) (via electroporation) was struck onto TSA cam and incubated at 44 °C for 16-20 h to select for a single-crossover event. Large colonies (candidates for having undergone a single recombination) were struck on TSA cam. Individual colonies were inoculated into 5 mL TSB without antibiotic and grown shaking at 250 rpm at 30 °C. Cultures were serially passaged twice daily for 5-10 cycles to promote a second recombination event. Following outgrowth, cultures were serially diluted and plated onto TSA containing 100 ng/mL anhydrotetracycline hydrochloride (Thermo Fisher Scientific #J66688MB) to enrich for double-crossover recombinants via counterselection (∼101-3 CFU plated corresponding to the 10-5-7 dilutions of bacterial suspension). Resulting colonies were patched onto TSA and TSA cam and PCR-verified to confirm loss of pKQ002 and successful allelic exchange.

### Vector construction for complementation at the *S. aureus* chromosomal Φ11 *attB* **site**

The integration plasmids pLL2787 and pLL39 were obtained from Addgene (pLL2787: plasmid #15460; pLL39: plasmid #15458) [68]. Plasmid pLL2787 was propagated in *E. coli* DH5α with amp selection, isolated, and electroporated into *S. aureus* RN4220 [104] with selection on TSA cam, following the same transformation and selection protocol as described for pKQ002 above.

For construction of complementation plasmids, pLL39 was linearized with SmaI (NEB R0141S). Complementation fragments were PCR-amplified from *S. aureus* JE2 using Phusion™ High-Fidelity Master Mix, as described for the *alr1* homology arms. **Table S5** lists primer sequences. After purification, PCR fragments were inserted into the SmaI site of pLL39 using a Gibson Assembly Kit (NEB E5510S). Assembly mixtures were transformed into chemically competent *E. coli* DH5α. Transformants were selected on LB agar supplemented with 50 μg/mL spectinomycin (spc) then colony purified thrice. Plasmid DNA was isolated using the QIAprep Spin Miniprep Kit (Qiagen 27104) and eluted in nuclease-free water, as described above. Purified plasmids were electroporated into *S. aureus* RN4220 carrying pLL2787, using the conditions described above for pKQ002 into *S. aureus*. pLL39 is nonreplicative in *S. aureus* and maintained only when integrated on the chromosome at the Φ11 attachment site (*attB*). Transformants were selected on TSA with 3 μg/ml tetracycline (tet) and colony-purified thrice prior to freezer stocking. Complementation constructs were transduced from RN4220 to each target *S. aureus* as described below. **Table S4** lists all strains and plasmids, and **Table S5** lists primers used to generate each complementation construct. The Φ11 *attB* site is at the beginning of SAUSA300_RS10150 and complementation constructs integrate at CCATG|GGAAG.

### Φ11 phage transduction of Tn insertions into specific genetic backgrounds

Tn insertion mutants in *dat*::Tn, *alr1*::Tn, *fmtA*::Tn, and *hla*::Tn from the NTML (BEI Resources NR-49947) [76] were transduced into our lab strain of *S. aureus* JE2 (KPL2115) via phage transduction as previously described [105]. Briefly, Φ11 lysates from the Tn-bearing strain of interest were generated by embedding 100 µL of mid-exponential phase (OD600 = 0.3-0.5) of the Tn mutant strain in 0.75% TSB soft agar with at least 108 PFUs of Φ11. After growing for 16-20 h at 37 °C, the soft agar was collected, and Φ11 (with some phage carrying the Tn insertion of interest) was purified and concentrated by centrifugation at 5,000-7,800 rpm with a 100 kDa MWCO Amicon® Ultra Centrifugal Filter (Millipore UFC9100). Final lysates were passed through 0.22 µm Millex™ polyethersulfone syringe filter (Millipore SLGP033N) to remove bacteria.

Each recipient strain (400 µL of mid-log bacterial growth in TSB to OD600 = 0.3-0.5) was infected with 400 µL (≥ 108 PFUs) Φ11 from the donor strain of interest (20 min, 37 °C, 250 rpm). Next, samples were immediately placed on ice and the infection was deactivated with 3 mL 20 mM sodium citrate (pH 5.0) (Sigma-Aldrich #S1804). Infected cells were pelleted at 6,000 x *g* for 10 min at 4 °C and washed twice with 5 mL of 20 mM sodium citrate (pH 5.0). The final pellet was resuspended in 200-300 µL of 20 mM sodium citrate and spread plated on antibiotic-selective TSA. Colonies displaying antibiotic resistance were colony-purified thrice and stocked as described above. PCR was performed using GoTaq (Promega, #M7123) using an insertion-specific primer (**Table S5**) and the upstream/buster primers [95] to confirm the presence of the Tn insertion.

### Preparation of electrocompetent bacteria

A protocol for preparing electrocompetent cells (*E. coli*, *S. aureus*) was adapted from the instruction manual of the BioRad Gene Pulser XCell™ Electroporation System (#1652660) and Zeden et al. [103], and we replaced sucrose with 10% glycerol. *E. coli* cultures were grown in LB broth (Difco™ 244620) shaking at 37 °C to an OD600 of 0.4-0.6. Cultures were chilled on ice for 10-15 min to stop growth, then bacteria were pelleted by centrifugation at 4 °C, 4000 x *g* for 10 min (Eppendorf 5430R) and the supernatant discarded. The pellet was gently resuspended in 50 mL ice-cold sterile 10% glycerol (Sigma Aldrich #G5516) and divided into two 25 mL aliquots in prechilled 50 mL conical tubes on ice, then bacteria were washed twice more with an equal volume 10% glycerol by centrifugation (4 °C, 4000 x *g*, 10 min) followed by resuspension in ice-cold 10% glycerol, for a total of three washes. After the final wash, pellets were resuspended in 0.5 mL ice-cold 10% glycerol and pooled to a final volume of 1 mL. Electrocompetent cells were aliquoted into prechilled microcentrifuge tubes (∼110 µL per tube) (Eppendorf 20901-551), flash-frozen in a dry ice-ethanol bath, and stored at −80 °C.

Overnight *S. aureus* cultures grown in TSB (3 mL) were diluted to an OD600 of 0.4-0.6 in 50 mL and incubated shaking (250 rpm) at 37 °C until reaching an OD600 of 0.8-0.9. We transferred 25 mL of culture each to two 50 mL conical tubes, chilled on ice for 10 min, then harvested cells by centrifugation (5000-7000 x *g*, 10 min, 4 °C). Supernatants were discarded and pellets were rinsed with 45 mL ice-cold sterile Milli-Q water without resuspension, then collected again via centrifugation. Each pellet was first resuspended in 45 mL ice-cold MilliQ water, followed by centrifugation, then resuspended in 10 mL ice-cold 10% glycerol, followed by two more washes with 10 mL ice-cold 10% glycerol. After the final wash, cells were resuspended in 2 mL ice-cold 10% glycerol and transferred to a 5 mL microcentrifuge tube. Cells were pelleted in a microcentrifuge (10,000 x *g*, 2 min, 4 °C), resuspended in 1 mL, pelleted again and resuspended in 250 µL ice-cold 10% glycerol, and maintained on ice throughout. Electrocompetent cells were aliquoted into prechilled tubes (∼110 µL per tube), flash-frozen in a dry ice-ethanol bath, and stored at −80 °C.

### Electroporation

Electroporation was performed using a BioRad Gene Pulser XCell™ (BioRad #1652660) with a prechilled 0.1 cm gap electroporation cuvette (Biorad #1652089). For transformation into *E. coli* according to [106], 110 µL of electrocompetent *E. coli* cells were combined with 500 ng plasmid DNA, transferred to a cuvette and electroporated at 200 Ω, 1.8 kV, and 25 µF. Immediately after, we added 0.9 mL of RT SOC outgrowth medium (NEB #B9020S) directly to the cuvette, and the cell suspension was transferred to microcentrifuge tubes and incubated at 37 °C for 1.5-2 h without shaking to allow recovery prior to inoculation onto antibiotic-selective LB agar medium followed by overnight growth to identify transformants. For transformation into *S. aureus* according to [103], 110 µL of electrocompetent *S. aureus* cells were combined with up to 5 µg plasmid DNA, transferred to a cuvette, and electroporated at 100 Ω, 2.5 kV, and 25 μF. Immediately after, 0.9 mL of RT TSB supplemented with 0.5 M D-sucrose was added directly to the cuvette and the cell suspension was transferred to microcentrifuge tubes. Cultures were incubated at 37 °C for 1.5-2 h without shaking to allow recovery then inoculated onto antibiotic-selective agar medium for overnight growth at 37 °C to identify transformants. TSA was used for *S. aureus* strains and LB agar was used for *E. coli* strains.

### Colony PCR

To enhance success with colony PCR of *S. aureus*, a single colony of interest was suspended in 100 µL of nuclease-free water with 2 µL lysostaphin (10 µg/ml) (Sigma Aldrich #L7386), incubated 15 min at 37 °C, heated to 95 °C for 10 min, frozen at 80 °C for ≥ 15 min, and thawed before each PCR reaction. If colony PCR was unsuccessful, genomic DNA was prepared for each colony of interest prior to the PCR assay.

### Whole Genome Sequencing of *dat*::Tn P*datdat*

High quality genomic DNA was prepared from KPL4549 (*hla*::Tn (spcR) *dat*::Tn (ermR) attB::pLL39::P*datdat*) as described above in DNA extraction for TnSeq. DNA was sequenced using Oxford Nanopore and mapped to the JE2 reference genome (USA300_FPR3757) as a fee for service from Plasmidsaurus.

### Antibiotic cassette exchange

The NTML Tn mutants all contain an erm-resistance gene *ermB* [76]. To generate *hla*::Tn with a spc-resistance cassette and both *dat*::Tn insertions with tet-resistance cassettes, antibiotic cassette exchange was performed as described in Bose et al. [95] with temperature-sensitive allelic exchange plasmids in *S. aureus* RN4220 that enabled the replacement of the erm-resistance gene in *bursa aurealis* with tet-, or spc-resistance genes. We used *S. aureus* RN4220 containing pSPC (#NRC-49934) or pTET (#NRC-49936) from the NTML Genetic Toolbox (BEI Resources #NR-49947). Plasmids pSPC and pTET are stable at 30 °C and confer resistance at a concentration of 1,000 µg/mL spc, and 5 µg/mL tet, respectively. pSPC or pTET were isolated and electroporated into the Tn mutant strain of interest as described above, and strains were confirmed via PCR to have the plasmid of interest (see **Table S5** for primers). The first and second recombination events were generated as described above for allelic exchange. Resulting colonies were patched onto TSA, TSA cam, TSA erm, and TSA spc or TSA tet to confirm loss of the plasmid, loss of the erm-resistance cassette, and introduction of the spc- or tet- resistance cassette. PCR was performed to confirm presence of the correct antibiotic-resistance cassette using primers listed in **Table S5**.

### Chemically defined medium (CDM)

We used a CDM with all 20 amino acids (CDM20) and a CDM lacking any alanine (CDM19-ala) as we previously described for nasal *Corynebacterium* species [72] without Tween80. The MOPS-buffered base CDM included a 2X solution of a Teknova medium prepared from Teknova 10X MOPS buffer (#M2101), Teknova 0.132 M potassium phosphate dibasic solution used to a final concentration (f.c.) of 2.64 mM (#M2102), 10x ACGU solution (#M2103), our 100x in-house vitamin stock (f.c. 1x) [72], plus 1,000x ferric chloride (f.c. 0.2 mM), and 400x lipoic acid (f.c. 0.0192 mM). The pH of the medium was adjusted to 7-7.5 before use. We prepared a 5X solution mixture of 20 amino acids in MilliQ water; the f.c. of each after the addition to base CDM was as follows: 1.6 mM L-alanine (omitted in CDM19-ala), 10.4 mM L-arginine, 0.8 mM L-asparagine, 0.8 mM L-aspartic acid, 0.2 mM L-cysteine, 1.2 mM L-glutamic acid, 1.2 mM L-glutamine, 1.6 mM L-glycine, 0.4 mM L-histidine, 0.8 mM L-isoleucine, 1.6 mM L-leucine, 0.8 mM L-lysine, 0.4 mM L-methionine, 0.8 mM L-phenylalanine, 0.8 mM L-proline, 20 mM L-serine, 0.8 mM L-threonine, 0.2 mM L-tryptophan, 0.4 mM L-tyrosine, and 1.2 mM L-valine.

### Growth curves in liquid media

Individual strains were struck onto TSA and incubated at 37 °C overnight for 16-20 h. A single colony from each plate was inoculated into 1 mL of CDM20 and grown overnight shaking at 250 rpm at 34 °C. Resulting cultures were adjusted to an OD600 equal to 0.5 in the corresponding medium. For each strain, 15 µL of the resuspension was added to 135 µL of TSB, CDM20, or CDM19-ala in a Corning Costar 96-well non-treated polystyrene flat bottom plate (Corning #3370). Growth was monitored in a BioTek Synergy H1 Multimode Reader at 34 °C with continuous orbital shaking for 24 h and the OD600 was recorded hourly.

### Competition experiments

The bacterial strains used in competition experiments were prepared for colonization as described above with a few modifications. Each strain was separately resuspended to an OD600 equal to 10^5^ CFUs in 7.5 µL EBSS. The initial inoculum for competition conditions was made by mixing an equal volume of each competing strain then 15 µL of the mix was inoculated onto the apical surface of each HNO-ALI, such that 105 CFUs of each strain was inoculated (for a total of 2 x 105 bacterial CFUs). For HNO-ALI collection and CFU enumeration, 20 µL drips were plated on both TSA and TSA with antibiotic to differentiate between total CFUs (TSA) and CFUs of the Tn(erm) or Tn(tet)-containing competitor. TSA erm 5 μg/ml was used for mutants in the JE2 and 502A strain background, and TSA tet 1 μg/ml was used for mutants in the AH2623 or KPL2715 strain background.

### Chemical complementation of bacterial mutants on HNO-ALI

Each bacterial strain of interest was inoculated onto HNO-ALI as described above. For chemical complementation experiments, 5 µL of the amino acid of interest or EBSS was inoculated directly following bacterial inoculation. To supplement *S. aureus* with exogenous amino acids, D-alanine (Millipore Sigma #A7377), L-alanine (Millipore Sigma #A7627), D-glutamic acid (Thermo Scientific Chemicals #A1419106), and L-glutamic acid (Avantor #207706) were resuspended in EBSS to 150 mM, 15 mM, or 1.5 mM, and the pH was adjusted to 7.0. Inoculation of 5 µL amino acid onto an HNO-ALI colonized with 15 µL *S. aureus* led to an estimated final concentration of amino acid supplementation of 37.5 mM, 3.75 mM, and 0.375 mM, respectively.

### Cocolonization of HNO-ALI with *C. pseudodiphtheriticum* and *S. aureus*

The bacterial strains used in cocolonization experiments were prepared for colonization as described above with a few modifications. *S. aureus* (parent or *dat*::Tn) were grown on TSA as described and separately resuspended to an OD600 equal to 10^5^ CFUs in 7.5 µL EBSS. *Corynebacterium pseudodiphtheriticum* KPL1989 [83] was struck for single colonies and grown atop a 0.2-μm polycarbonate membrane on TSA for 48 h at 34 °C in a 5% CO2-enriched incubator. The day of colonization, 5-10 colonies were resuspended in 5 ml EBSS, spun at 10,000 x *g* for 10 min at RT, and the cell pellet was resuspended in 1 mL EBSS. The bacterial resuspension was then normalized to an OD600 = 10^6^ CFUs in 7.5 µL EBSS. Each resuspension was then mixed in equal volume to yield a 15 µL inoculum containing 105 CFUs *S. aureus* (either parent or *dat*::Tn) and 106 CFUs *C. pseudodiphtheriticum*. HNOs were colonized with each condition for 24 h at 34 °C in a 5% CO2-enriched incubator.

### Statistical Analysis

All data processing, statistical analyses, and visualization were performed using R [107] within the RStudio integrated development environment [108]. Reproducibility was ensured through the use of a controlled package environment managed with the renv package [109]. A complete list of all R packages used, including version numbers and session information, as well as instructions for reproducing the computational environment using renv are available at: https://klemonlab.github.io/DatCol_Manuscript/RSession.html.

Statistical analyses were performed using a standardized linear mixed-effects modeling (LMM) workflow implemented via custom functions and described in detail at: https://klemonlab.github.io/DatCol_Manuscript/Manuscript_DatCol_LMMStats.html.

LMMs were fit using the lme4 package [110], with fixed effects specified according to the experimental design and random effects included to account for structured sources of variability. Specifically, experimental date (i.e., independent experiment) was modeled as a random effect nested within HNO donor line when applicable, to control for batch effects and biological variation. When singular fits were detected, or when only one HNO donor line was used, the random-effects structure was simplified to only include experimental date.

Log transformation of the response variable was applied when appropriate to meet model assumptions. For growth curves in **Fig. 3A** and **Fig. S3A-C**, the area under the curve (AUC) was computed using the trapezoidal rule from the pracma package. Statistical significance of fixed effects was assessed using analysis of variance (ANOVA) with denominator degrees of freedom estimated using the Satterthwaite approximation, as implemented in the lmerTest package [111]. Post hoc pairwise comparisons were conducted using estimated marginal means computed with the emmeans package [112]. In Figures 2 and 5, we used a one sample *t*-test to compare the competition index (CI) of each mutant to the theoretical parental-strain value of 1. *P*-values were adjusted for multiple comparisons using the Holm method [113].

Model-derived estimated marginal means, contrasts, and confidence intervals were extracted and, where applicable, back-transformed to the original response scale for interpretability and visualization (**Table S2**). Complete analysis code for each dataset, including specific model formulations, is available at https://klemonlab.github.io/DatCol_Manuscript/.

## Data Availability

All dataframes used in this study and all R code used to analyze data and generate figures are available at https://klemonlab.github.io/DatCol_Manuscript/

## Acknowledgments

We thank the donors who make HNO-ALI possible. We thank Anthony Richardson for sharing his *S. aureus* LAC Tn library, Jennifer Walker and Jesus Duran Ramirez for the gift of transducing phage and help with transduction protocols, The NTML (NR-48501) and plasmids from the NTML Toolkit were provided by B.E.I. (http://www.beiresources.org/).

This research was supported in part by funding from the National Institute of Allergy and Infectious Diseases of the National Institutes for Health (NIH) under award U19AI157981 (K.P.L., S.E.B.) and award F31AI172324 (A.I.B.), the National Institute of General Medical Sciences (NIGMS) of the NIH under award R35GM160019 (C.B.I.), Baylor College of Medicine (via seed funds to K.P.L.), and the University of Oklahoma (via start-up funds to C.B.I.).

## Author Contributions

Conceptualization: A.I.B., C.B.I., K.P.L. Methodology: A.I.B., K.A.Q., I.F.E., L.A.K., C.B.I., K.P.L. Investigation: A.I.B., K.A.Q. Resources: I.F.E., X.L.Z., S.E.B., C.B.I. Data curation: A.I.B., I.F.E., C.B.I. Formal analysis: A.I.B., I.F.E. Software: I.F.E. Validation: A.I.B, K.A.Q. Visualization: A.I.B., I.F.E. Writing – original draft: A.I.B., K.A.Q., I.F.E., M.A.L., K.P.L. Writing – review & editing: A.I.B., K.A.Q., I.F.E., M.A.L., L.A.K., X.L.Z., S.E.B., C.B.I., K.P.L. Funding acquisition: A.I.B., S.E.B., C.B.I., K.P.L. Supervision: I.F.E., K.P.L.

## Supporting Materials

**Figure S1.**
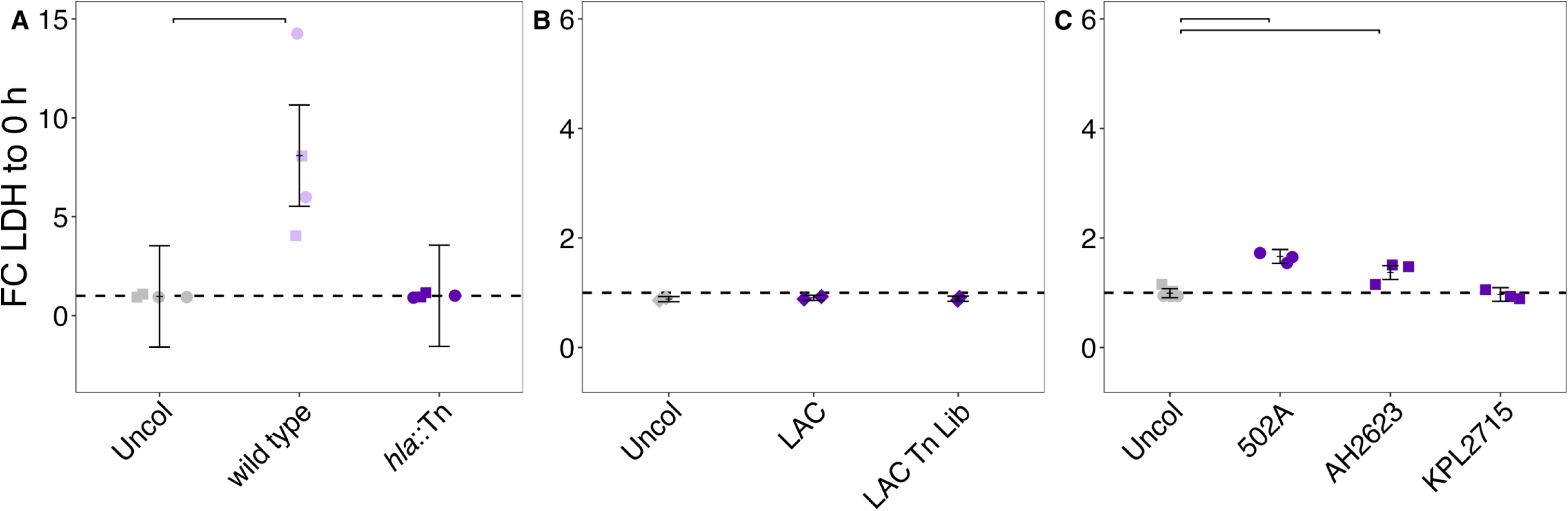
Cytotoxicity profiling of *S. aureus* strains on HNO–ALI. (**A**) *S. aureus* LAC induced cytotoxicity by 24 h in an *hla*-dependent manner in HNO-ALI generated using a recently revised protocol that has improved on-time production. (**B**) No cytotoxicity was caused by LAC (CC8) or LAC Tn library after 8 h colonization in current HNO-ALI. (**C**) There was minimal HNO-ALI cytotoxicity caused by 502A (CC30) and AH2623 (CC5) by 16 h of colonization and no apparent toxicity caused by KPL2715 (CC30) after 24 h colonization. LDH was quantified as a fold change from the final timepoint of colonization compared to the amount of LDH detected at 1 hour before colonization. *n* ≥ 2 independent experiments in a minimum of 1 HNO line (HNO204, circles; HNO918, squares; or HNO919, diamonds). Fold change between groups was compared using an LMM with colonizing strain as fixed effect and experimental date as a random effect, with contrasts performed relative to the uncolonized condition and adjusted for multiple comparisons using the Holm method. Horizontal brackets represent statistically significant comparisons *p* < 0.05. Vertical brackets represent the model-predicted mean values and confidence intervals ± 2 x SEM. Statistical analysis is in **Table S2E**.

**Figure S2.**
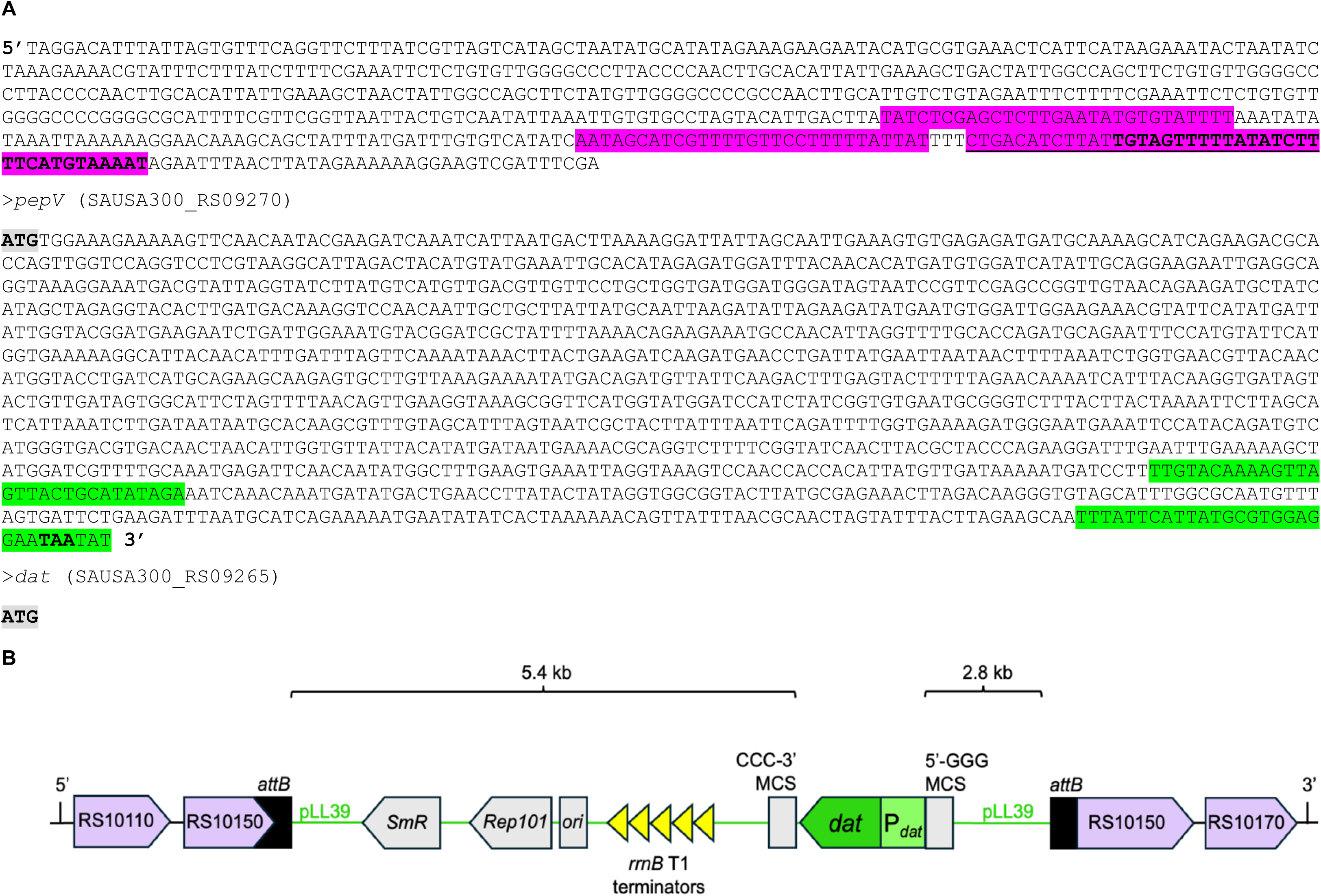
P*dat* is the only promoter likely to promote *dat* transcription in KPL4549 (*dat*::Tn P*datdat)*. **(A)** SAPPHIRE-predicted promoter sequences identified 5’ of and within the *pepV* coding region of *S. aureus* for complementation construction of P*_pepV_* (magenta highlight) and P*_dat_* (green highlight), respectively. (Overlapping sequences are underlined and bolded for differentiation). *pepV* and *dat* start codons are highlighted in gray. **(B)** Schematic representation of single-copy chromosomal complementation of *dat* from whole-genome-sequenced KPL4549 (*hla*::Tn (spc^R^) *dat*::Tn (erm^R^) *attB*::pLL39::P*_dat_dat*). The Φ11 *attB* integration site is located 46 bp into the coding region of SAUSA300_RS10150. Recombination with the pLL39-derived complementation construct pAB004 (*P_dat_dat*) occurred at 5′-CCATG|GGAAG-3’. The complementation construct (*P_dat_dat*) was assembled at *SmaI* (5’-GGGCCC-3’) in the multiple cloning site (MCS) by Gibson assembly into pLL39 (see **Table S5** for complementation oligomers). Relevant plasmid elements include the spectinomycin resistance cassette (*SmR*), *Rep101* and SC101 origin in *E. coli*, and *rrnB* T1 transcriptional terminators to minimize transcriptional readthrough. The *P_dat_dat* complementation insert is in the opposite orientation of the surrounding chromosomal genes, as shown on the displayed 5′ → 3′ strand.

**Figure S3.**
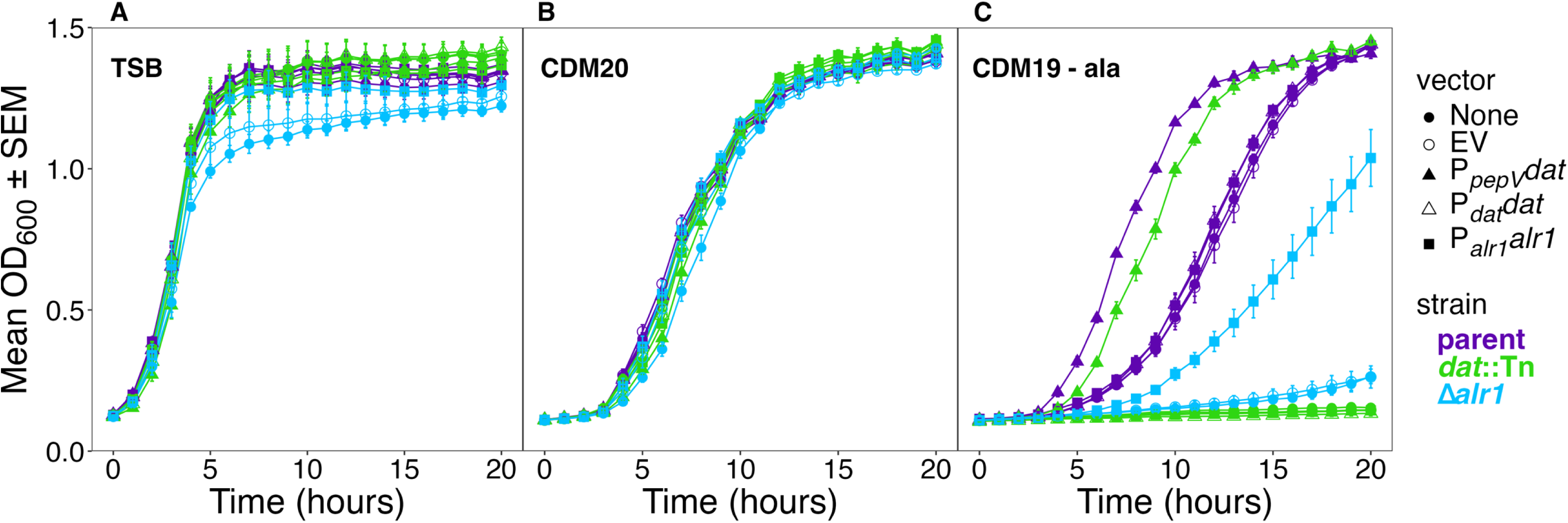
Complete set of growth curves for all *S. aureus* strains (mutants and their complements) in the KPL4530 background. Panels correspond to (**A**) TSB, (**B**) CDM20, and (**C**) CDM19-alanine liquid growth media. The parental *S. aureus* KPL4530 (JE2 *hla*::Tn(spc)) (purple) and isogenic *dat*::Tn(erm) (green) and Δ*alr1* (blue) mutants each carried either no vector (filled circles), empty vector (EV) pLL39 (empty circles), P*_pepV_dat* (filled triangles), P*_dat_dat* (empty triangles), or P*_alr1_alr1* (filled squares), inserted at the Φ11 *attB* site. *n* = 3 independent experiments. Each data point is the average OD600 ± SEM of all 3 experiments at that time point. To assess differences in growth curves within each medium, we compared the area under the curve (AUC) using an LMM with strain as a fixed effect and experimental date as a random effect. We performed pairwise contrasts comparing the parental strain carrying either the empty vector (EV) or no vector (None) to all other strains. In addition, we evaluated contrasts among plasmid variants within each mutant background (*dat*::Tn and *Δalr1*). Selected comparisons were adjusted for multiple testing using the Holm method, with statistical significance defined as *p* < 0.05. Statistical analysis is in **Table S2B**.

**Figure S4.**
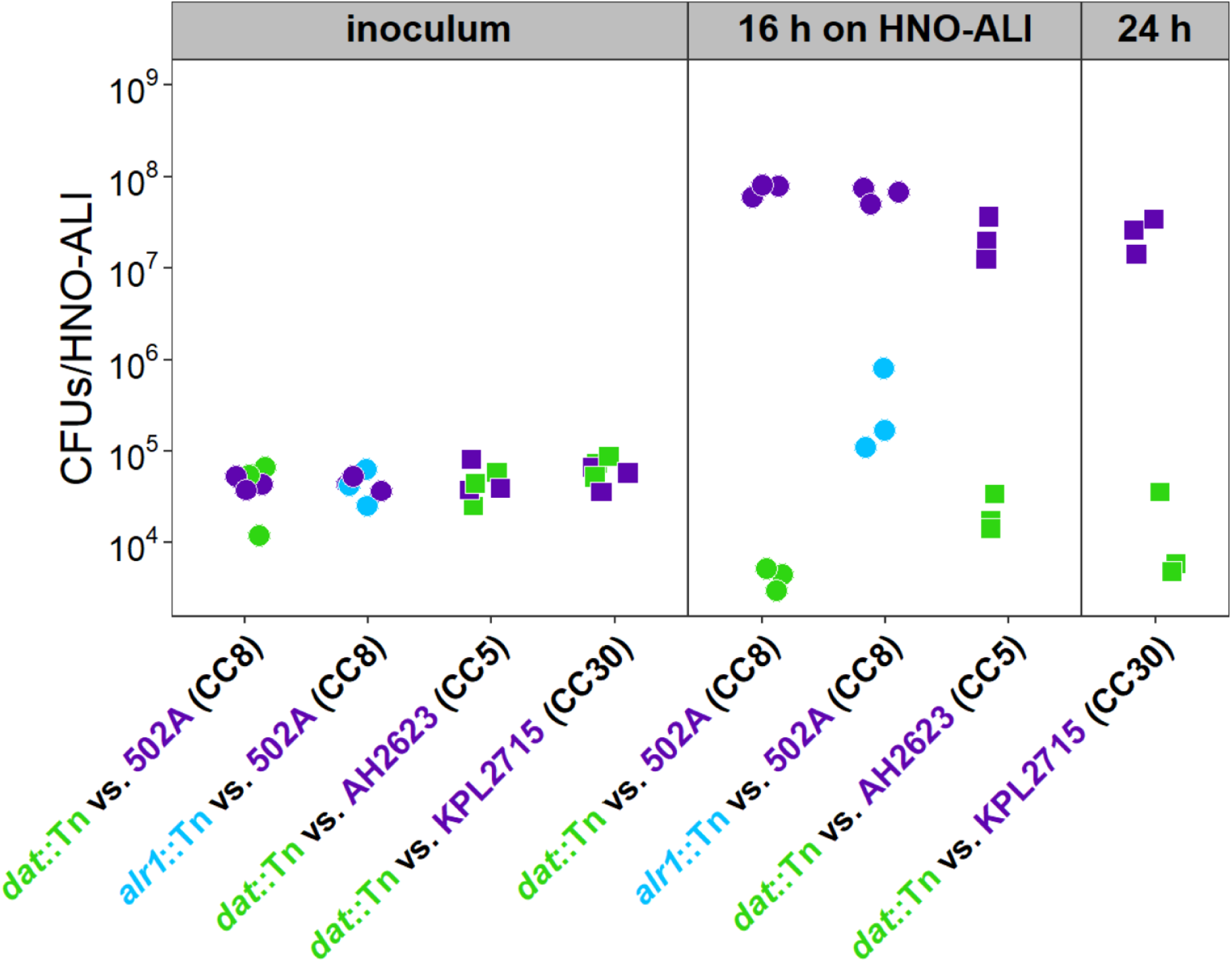
*dat* contributes to *S. aureus* fitness during nasal colonization across multiple clonal complexes. CFUs/HNO-ALI from *S. aureus* strains in Clonal Complex (CC) 8, 5, and 30. The parental strain (purple) outcompeted an isogenic mutant with a Tn disruption in either *dat* (green), or *alr1* (blue), respectively. HNO-ALI were cocolonized with 10^5^ CFUS/HNO-ALI each of the parental *S. aureus* strains, (502A – CC8, AH2623 – CC5, and KPL2715 – CC30) and each mutant of interest for 16 or 24 h at 34 °C in *n* = 3 independent experiments in either of 2 HNO lines (HNO204, circles or HNO918, squares).

**Table S1. TnSeq analysis (as a separate xlsx file)**

**• S1A. TnSeq Gene Level Analysis**

**• S1B. STRING Enrichment Analysis**

**Table S2. Statistical analyses (as a separate xlsx file)**

**• S2A. HNO-ALI CFU Competition Analysis**

**• S2B. Growth Curves OD600**

**• S2C. HNO-ALI CFU Complementation Analysis**

**• S2D. S. aureus coculture with C. pseudodiphtheriticum**

**• S2E. Cytotoxicity - LDH**

**Table S3.**
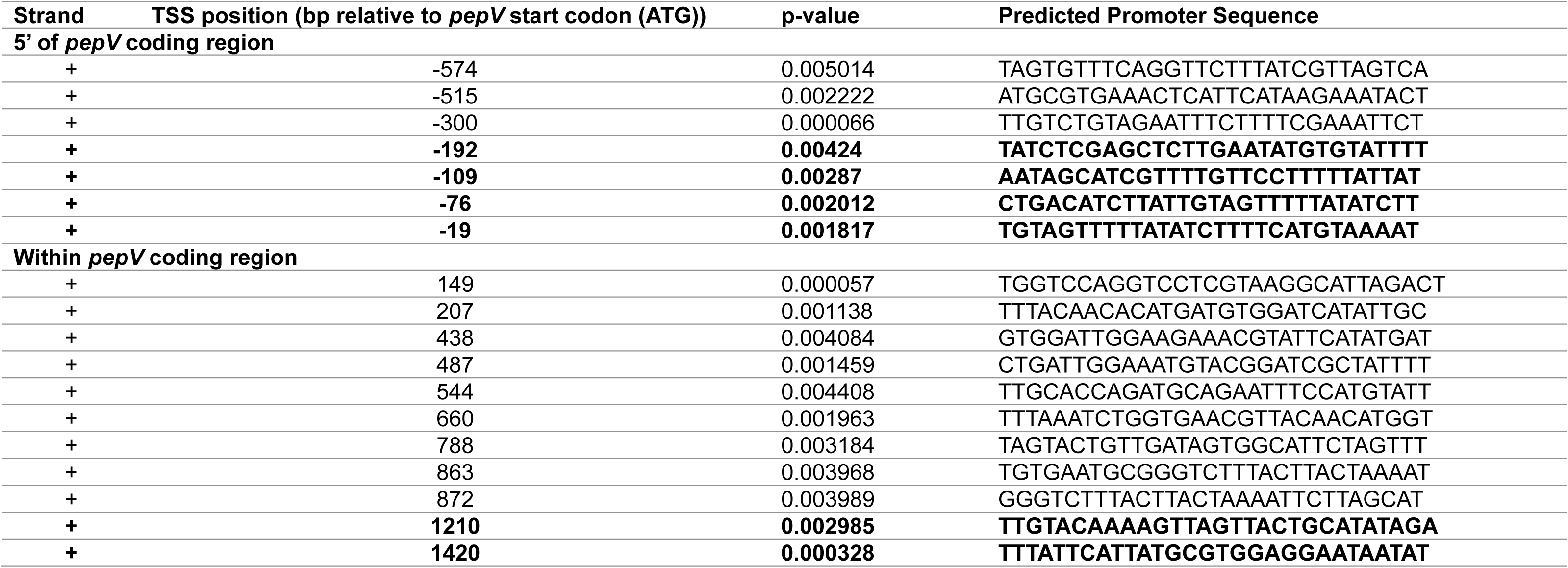
P*_dat_* and P*_pepV_* region SAPPHIRE promoter prediction results.

**Table S4.** Strains and plasmids used in this study.

| Species | Strain | Int. Ref. | Characteristics | Reference |
| --- | --- | --- | --- | --- |
| *E. coli* | NEB 5-alpha (#C2987) | KPL4489 | High-efficiency DH5α-derived cloning strain (endA<sup>−</sup>, T1 phage resistant) used for plasmid construction, Gibson assembly, and propagation | [1] |
| *E. coli* | NEB 5-alpha | KPL4565 | NEB 5-alpha transformed with Gibson assembled pKQ002 (*Δalr1*) | This study |
| *E. coli* | NEB 5-alpha | KPL4534 | NEB 5-alpha transformed with Gibson assembled pAB001 (pLL39::*P<sub>alr1</sub>alr1*) | This study |
| *E. coli* | NEB 5-alpha | KPL4535 | NEB 5-alpha transformed with Gibson assembled pAB004 (pLL39::*P<sub>dat</sub>dat*) | This study |
| *E. coli* | NEB 5-alpha | KPL4536 | NEB 5-alpha transformed with Gibson assembled pAB009 (pLL39::*P<sub>pepV</sub>dat*) | This study |
| *E. coli* | DC10b | KPL2100 | dam<sup>+</sup> dcm<sup>−</sup> derivative of DH10B, intermediate cloning host used prior to transformation into *S. aureus* | [2] |
| *E. coli* | DC10b | KPL4566 | DC10b (KPL2100) transformed with isolated pKQ002 (*Δalr1*) from NEB 5-alpha (KPL4565) | This study |
| *S. aureus* | LAC | KPL4403 | USA300 CA-MRSA strain LAC (MLST ST8, *spa* Type I, *SCCmec* Type IV, *agr* Type I) | [3] |
| *S. aureus* | LAC | KPL4447 | USA300 MRSA LAC saturated transposon library, with 77,161 independent insertions in genome | [4] |
| *S. aureus* | JE2 | KPL4465 | *hla*::Tn *bursa aurealis* transposon mutant (SAUSA300_RS05720<sup>1</sup>) in JE2 from the Nebraska Transposon Mutant Library (NTML) | [5] |
| *S. aureus* | JE2 | KPL4471 | NTML *dat*::Tn *bursa aurealis* transposon mutant (SAUSA300_RS09265) | [5] |
| *S. aureus* | JE2 | KPL4472 | NTML *alr1*::Tn *bursa aurealis* transposon mutant (SAUSA300_RS11155) | [5] |
| *S. aureus* | JE2 | KPL4474 | NTML *fmtA*::Tn *bursa aurealis* transposon mutant (SAUSA300_RS05155) | [5] |
| *S. aureus* | RN4220 | KPL4454 | Restriction-deficient (*hsdRMS*<sup>−</sup>) laboratory strain | [6] |
| *S. aureus* | RN4220 | KPL4537 | RN4220 transformed with pLL39 empty vector | This study |
| *S. aureus* | RN4220 | KPL4538 | RN4220 transformed with pAB004 (pLL39::*P<sub>dat</sub>dat*) from NEB 5-alpha (KPL4535) | This study |
| *S. aureus* | RN4220 | KPL4539 | RN4220 transformed with pAB009 (pLL39::*P<sub>pepV</sub>dat*) from NEB 5-alpha (KPL3536) | This study |
| *S. aureus* | RN4220 | KPL4540 | RN4220 transformed with pAB001 (pLL39::*P<sub>alr1</sub>alr1*) from NEB 5-alpha (KPL4534) | This study |
| *S. aureus* | RN4220 | KPL4542 | ***dat*::Tn (erm<sup>R</sup>)**, transduced (Φ11) from NTML *dat*::Tn (SAUSA300_RS09265) into *S. aureus* RN4220 | This study |
| *S. aureus* | RN4220 | KPL4546 | ***alr1*::Tn (erm<sup>R</sup>)** transduced (Φ11) from NTML *alr1*::Tn (SAUSA300_RS11155) into *S. aureus* RN4220 | This study |
| *S. aureus* | JE2 | KPL2115 | Plasmid-free derivative of USA300 MRSA strain LAC | [5] |
| *S. aureus* | JE2 | KPL4515 | ***dat*::Tn (erm<sup>R</sup>)**, transduced (Φ11) from NTML *dat*::Tn (SAUSA300_RS09265) into *S. aureus* JE2 | This study |
| *S. aureus* | JE2 | KPL4516 | ***hla*::Tn (erm<sup>R</sup>)**, transduced (Φ11) from NTML *hla*::Tn (SAUSA300_RS05720) (KPL4465) into JE2 | This study |
| *S. aureus* | JE2 | KPL4530 | ***hla*::Tn (spc<sup>R</sup>)**, transduced (Φ11) from NTML *hla*::Tn (SAUSA300_RS05720) into JE2, followed by antibiotic cassette exchange via pSPC (erm<sup>R</sup> → spc<sup>R</sup>) | This study |
| *S. aureus* | JE2 | KPL4532 | ***Δalr1***, *S. aureus* JE2 transformed with isolated pKQ002 in *E. coli* DC10b (KPL4566) and allelic exchange performed for markerless deletion of *alr1* (See Methods) | This study |
| *S. aureus* | JE2 | KPL4541 | ***hla*::Tn (spc<sup>R</sup>) *Δalr1***, transduced (Φ11) from *hla*::Tn (spc<sup>R</sup>) into JE2 *Δalr1* (KPL4532) | This study |
| *S. aureus* | JE2 | KPL4545 | ***hla*::Tn (spc<sup>R</sup>) *dat*::Tn (erm<sup>R</sup>)**, transduced (Φ11) from *hla*::Tn (spc<sup>R</sup>, KPL4530) into *dat*::Tn (erm<sup>R</sup>, KPL4515) | This study |
| *S. aureus* | JE2 | KPL4556 | ***hla*::Tn (spc<sup>R</sup>) *fmtA*::Tn (erm<sup>R</sup>)**, transduced (Φ11) from *fmtA*::Tn (erm<sup>R</sup>, KPL4474) into *hla*::Tn (spc<sup>R</sup>, KPL4530) | This study |
| *S. aureus* | JE2 | KPL4553 | ***hla*::Tn (spc<sup>R</sup>) attB::pLL39**, KPL4530 with the pLL39 empty vector integrated at the Φ11 *attB* site | This study |
| *S. aureus* | JE2 | KPL4543 | ***hla*::Tn (spc<sup>R</sup>) attB::pLL39::*P<sub>dat</sub>dat***, KPL4530 with pAB004 (pLL39::*P<sub>dat</sub>dat*) integrated at the Φ11 *attB* site | This study |
| *S. aureus* | JE2 | KPL4552 | ***hla*::Tn (spc<sup>R</sup>) attB::pLL39::*P<sub>pepV</sub>dat***, KPL4530 with pAB009 (pLL39::*P<sub>pepV</sub>dat*) integrated at the Φ11 *attB* site | This study |
| *S. aureus* | JE2 | KPL4544 | ***hla*::Tn (spc<sup>R</sup>) attB::pLL39::*P<sub>alr1</sub>alr1***, KPL4530 with pAB001 (pLL39::*P<sub>alr1</sub>alr1*) integrated at the Φ11 *attB* site | This study |
| *S. aureus* | JE2 | KPL4547 | ***hla*::Tn (spc<sup>R</sup>) *dat*::Tn (erm<sup>R</sup>) attB::pLL39**, KPL4545 with the pLL39 empty vector integrated at the Φ11 *attB* site | This study |
| *S. aureus* | JE2 | KPL4549 | ***hla*::Tn (spc<sup>R</sup>) *dat*::Tn (erm<sup>R</sup>) attB::pLL39::*P<sub>dat</sub>dat***, KPL4545 with pAB004 (pLL39::*P<sub>dat</sub>dat*) integrated at the Φ11 *attB* site | This study |
| *S. aureus* | JE2 | KPL4550 | ***hla*::Tn (spc<sup>R</sup>) *dat*::Tn (erm<sup>R</sup>) attB::pLL39::*P<sub>pepV</sub>dat***, KPL4545 with pAB009 (pLL39::*P<sub>pepV</sub>dat*) integrated at the Φ11 *attB* site | This study |
| *S. aureus* | JE2 | KPL4548 | ***hla*::Tn (spc<sup>R</sup>) *dat*::Tn (erm<sup>R</sup>) attB::pLL39::*P<sub>alr1</sub>alr1***, KPL4545 with pAB001 (pLL39::*P<sub>alr1</sub>alr1*) integrated at the Φ11 *attB* site (extra copy of *alr1*) | This study |
| *S. aureus* | JE2 | KPL4554 | ***hla*::Tn (spc<sup>R</sup>) *Δalr1* attB::pLL39**, *hla*::Tn (spc<sup>R</sup>) *Δalr1* (KPL4541) with pLL39 empty vector integrated at the Φ11 *attB* site | This study |
| *S. aureus* | JE2 | KPL4551 | ***hla*::Tn (spc<sup>R</sup>) *Δalr1* attB::pLL39::*P<sub>alr1</sub>alr1***, KPL4541 with pAB001 (pLL39::*P<sub>alr1</sub>alr1*) integrated at the Φ11 *attB* site | This study |
| *S. aureus* | JE2 | KPL4555 | ***hla*::Tn (spc<sup>R</sup>) *alr1*::Tn (erm<sup>R</sup>)**, transduced (Φ11) from NTML *alr1*::Tn (erm<sup>R</sup>, KPL4472) into *hla*::Tn (spc<sup>R</sup>, KPL4530) | This study |
| *S. aureus* | JE2 | KPL4560 | ***dat*::Tn (tet<sup>R</sup>)**, generated via antibiotic cassette exchange (erm<sup>R</sup> → tet<sup>R</sup>) from *dat*::Tn (erm<sup>R</sup>, KPL4515) | This study |
| *S. aureus* | RN4220 | KPL4561 | ***dat*::Tn (tet<sup>R</sup>)**, transduced (Φ11) from *dat*::Tn (tet<sup>R</sup>, KPL4560) into RN4220 (KPL4454) | This study |
| *S. aureus* | JE2 | KPL4562 | ***dat*::Tn (tet<sup>R</sup>) *hla*::Tn (spc<sup>R</sup>)**, transduced (Φ11) from *hla*::Tn (spc<sup>R</sup>, KPL4530) into *dat*::Tn (tet<sup>R</sup>, KPL4560) | This study |
| *S. aureus* | 502A | KPL4558 | CC8 clinical isolate 502A once used for bacterial interference of invasive *S. aureus* infections (1963) | [7] |
| *S. aureus* | 502A | KPL4567 | ***dat*::Tn (erm<sup>R</sup>)**, transduced (Φ11) from *dat*::Tn (erm<sup>R</sup>) in RN4220 (KPL4542) into *S. aureus* 502A (KPL4458) | This study |
| *S. aureus* | 502A | KPL4567 | ***alr1*::Tn (erm<sup>R</sup>)**, transduced (Φ11) from *alr1*::Tn (erm<sup>R</sup>) in RN4220 (KPL4546) into *S. aureus* 502A (KPL4458) | This study |
| *S. aureus* | AH2623 | KPL4559 | Nasal Isolate susceptible to 80α phage, USA100 Blood Isolate #209 | [8] |
| *S. aureus* | AH2623 | KPL4563 | ***dat*::Tn (tet<sup>R</sup>)**, transduced (Φ11) from *dat*::Tn (tet<sup>R</sup>) (KPL4560) into *S. aureus* AH2623 (KPL4459) | This study |
| *S. aureus* | MNM000212 | KPL2715 | Primary nasal isolate (*mecA*<sup>+</sup>, CC30, ST30) per PubMLST | This study |
| *S. aureus* | KPL2715 | KPL4360 | Passage of KPL2715 sent for WGS (Microbial Genome Sequencing Center) | This study |
| *S. aureus* | KPL2715 | KPL4564 | ***dat*::Tn (tet<sup>R</sup>)**, transduced (Φ11) from *dat*::Tn (tet<sup>R</sup>) (KPL4561) into *S. aureus* nasal isolate (KPL4360) | This study |
| *Corynebacterium pseudodiphtheriticum* | KPL1989 | KPL1989 | Primary adult human nostril isolate | [9] |
| Plasmid | Host Strain | Int. Ref. | Characteristics | Reference |
| pJB38 | NE3001 (*E. coli* DH5α) |  | pUC19-based allelic exchange vector in *E. coli* and *S. aureus* (*Eco*<sup>2</sup> amp<sup>R</sup>; *Sau*<sup>3</sup> cam<sup>R</sup>) | [10] |
| pSPC | NE3003 (*S. aureus* RN4220) | KPL4521 | Allelic exchange plasmid containing homologous DNA from *bursa aurealis* with *aad9* (spectinomycin resistance) for exchange with *ermB* in *bursa aurealis* (*Eco* amp<sup>R</sup>; *Sau* cam<sup>R</sup> spc<sup>R</sup>) | [10] |
| pTET | NE3005 (*S. aureus* RN4220) | KPL4522 | Allelic exchange plasmid containing homologous DNA from *bursa aurealis* with *tetM* (tetracycline resistance) for exchange with *ermB* in *bursa aurealis* (*Eco* amp<sup>R</sup>; *Sau* cam<sup>R</sup> tet<sup>R</sup>) | [10] |
| pLL39 (#15458, Addgene) | *E. coli* | KPL4528 | *S. aureus*-*E. coli* shuttle vector (with tandem terminators at MCS) for single-copy integration at L54a *attB* and phi-11 *attB* sites (*Sau* spc<sup>R</sup>) | [11] |
| pLL2787 (#15460, Addgene) | *E. coli* | KPL4529 | Bacterial expression vector, encoding Φ11 integrase for single-copy integration at *attB* sites (*Eco* amp<sup>R</sup>) | [11] |
| pKQ002 | *S. aureus* JE2 | KPL4532 | pJB38-derived allelic exchange plasmid assembled by Gibson assembly containing ~700–900 bp flanking regions of *alr1* for markerless in-frame deletion in JE2 (*Eco* amp<sup>R</sup>; *Sau* cam<sup>R</sup>) | This study |
| pAB004 | *S. aureus* RN4220 | KPL4538 | pLL39-derived vector for complementation, containing *dat* under its putative 5' intraoperon promoter (*P<sub>dat</sub>dat*) (*Sau* spc<sup>R</sup>) | This study |
| pAB009 | *S. aureus* RN4220 | KPL4539 | pLL39-derived vector for complementation, containing *dat* under the *pepV* promoter (*P<sub>pepV</sub>dat*) in operon (*Sau* spc<sup>R</sup>) | This study |
| pAB001 | *S. aureus* RN4220 | KPL4540 | pLL39-derived vector for complementation, containing *alr1* under its native promoter (*P<sub>alr1</sub>alr1*) (*Sau* spc<sup>R</sup>) | This study |

<sup>1</sup> Locus tags correspond to *Staphylococcus aureus* gene annotations in AureoWiki.
<sup>2</sup> *Eco* = *E. coli*
<sup>3</sup> *Sau* = *S. aureus*

**Table S5.**
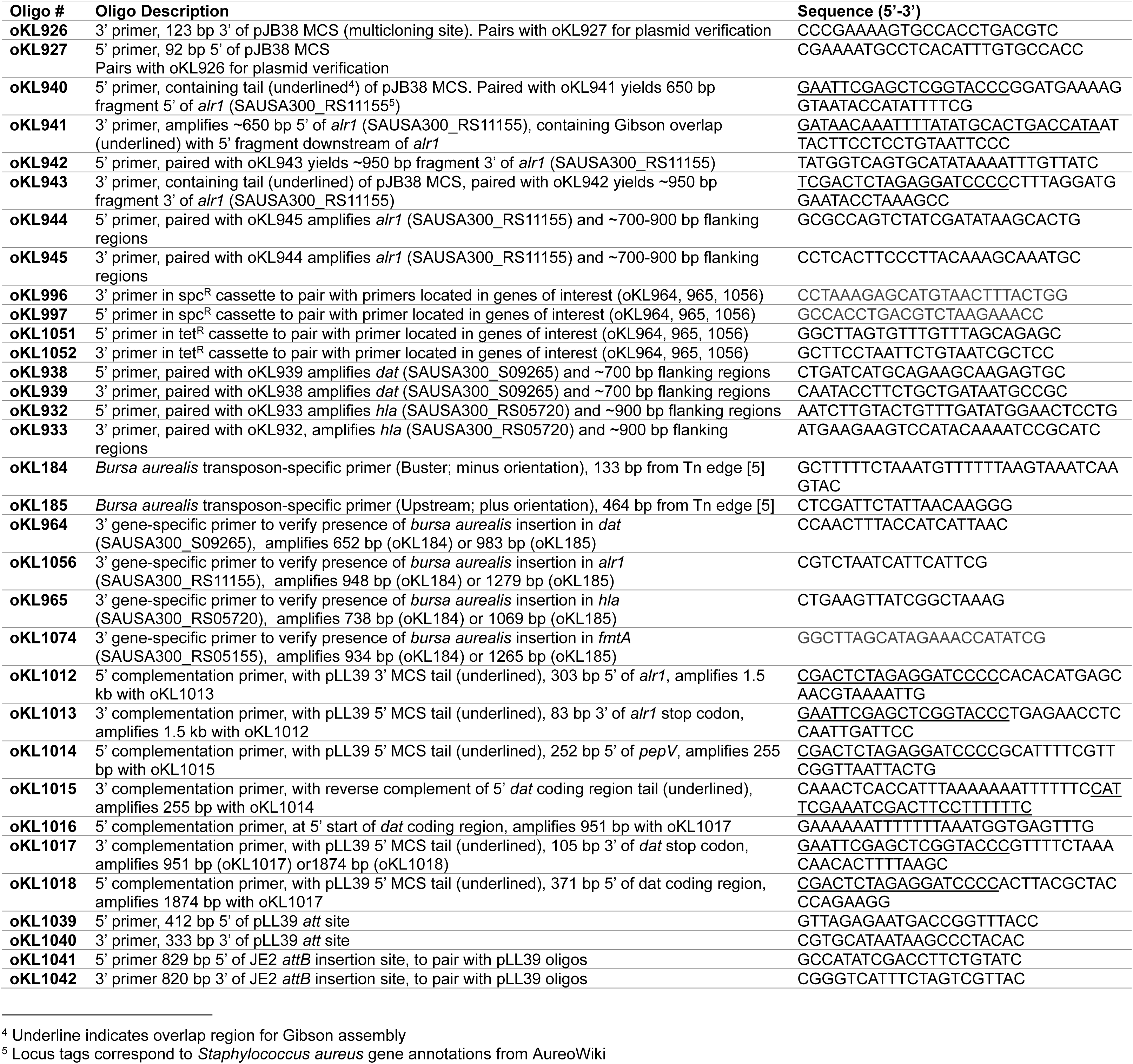
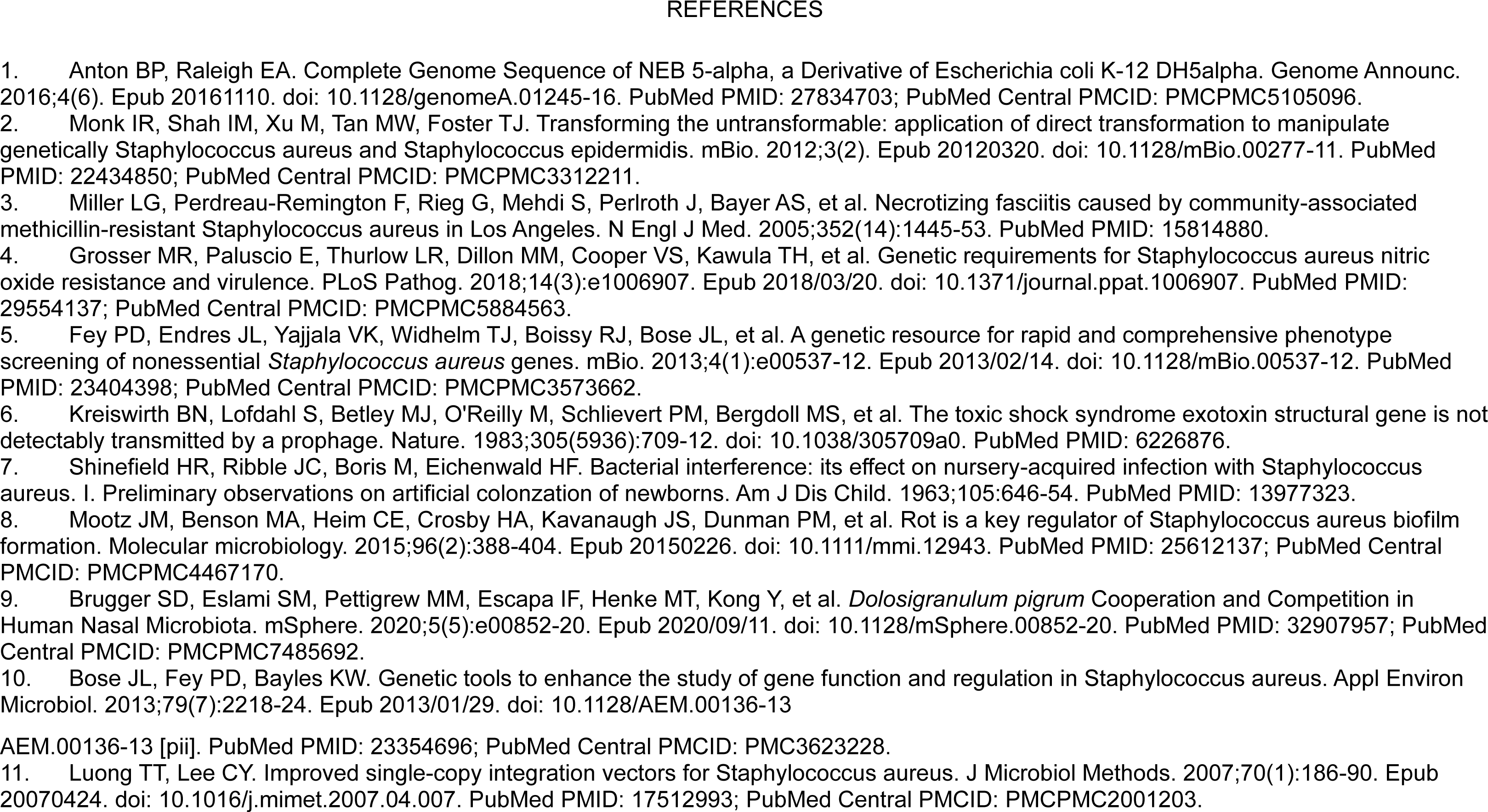
Oligonucleotides used in this study.

## REFERENCES

1. Collaborators GBDAR. Global mortality associated with 33 bacterial pathogens in 2019: a systematic analysis for the Global Burden of Disease Study 2019. Lancet. 2022;400(10369):2221–48. Epub 2022/11/25. doi: 10.1016/S0140-6736(22)02185-7. PubMed PMID: 36423648; PubMed Central PMCID: PMCPMC9763654.

2. von Eiff C, Becker K, Machka K, Stammer H, Peters G, Group FtS. Nasal carriage as a source of *Staphylococcus aureus* bacteremia. N Engl J Med. 2001;344(1):11–6. PubMed PMID: 11136954.

3. Wertheim HF, Vos MC, Ott A, van Belkum A, Voss A, Kluytmans JA, et al. Risk and outcome of nosocomial *Staphylococcus aureus* bacteraemia in nasal carriers versus non-carriers. Lancet. 2004;364(9435):703–5. Epub 2004/08/25. doi: 10.1016/S0140-6736(04)16897-9. PubMed PMID: 15325835.

4. Young BC, Wu CH, Gordon NC, Cole K, Price JR, Liu E, et al. Severe infections emerge from commensal bacteria by adaptive evolution. Elife. 2017;6. Epub 2017/12/20. doi: 10.7554/eLife.30637. PubMed PMID: 29256859; PubMed Central PMCID: PMCPMC5736351.

5. van Rijen M, Bonten M, Wenzel R, Kluytmans J. Mupirocin ointment for preventing *Staphylococcus aureus* infections in nasal carriers. Cochrane Database Syst Rev. 2008;(4):CD006216. Epub 2008/10/10. doi: 10.1002/14651858.CD006216.pub2. PubMed PMID: 18843708.

6. Bode LG, Kluytmans JA, Wertheim HF, Bogaers D, Vandenbroucke-Grauls CM, Roosendaal R, et al. Preventing surgical-site infections in nasal carriers of *Staphylococcus aureus*. N Engl J Med. 2010;362(1):9–17. Epub 2010/01/08. doi: 10.1056/NEJMoa0808939. PubMed PMID: 20054045.

7. Huang SS, Septimus E, Kleinman K, Moody J, Hickok J, Avery TR, et al. Targeted versus universal decolonization to prevent ICU infection. N Engl J Med. 2013;368(24):2255–65. Epub 2013/05/31. doi: 10.1056/NEJMoa1207290. PubMed PMID: 23718152; PubMed Central PMCID: PMCPMC10853913.

8. Nair R, Perencevich EN, Blevins AE, Goto M, Nelson RE, Schweizer ML. Clinical Effectiveness of Mupirocin for Preventing Staphylococcus aureus Infections in Nonsurgical Settings: A Meta-analysis. Clin Infect Dis. 2016;62(5):618–30. Epub 2015/10/28. doi: 10.1093/cid/civ901. PubMed PMID: 26503378.

9. Fritz SA, Bubeck Wardenburg J. A path forward for *Staphylococcus aureus* vaccine development. J Exp Med. 2024;221(10). Epub 2024/08/16. doi: 10.1084/jem.20240002. PubMed PMID: 39150449; PubMed Central PMCID: PMCPMC11329773.

10. WHO. Evidence-Based Recommendations on Measures for the Prevention of Surgical Site Infection: Preoperative Measures. Global Guidelines for the Prevention of Surgical Site Infection. WHO Guidelines Approved by the Guidelines Review Committee. 2nd ed. Geneva: World Health Organization; 2018.

11. CDC. Strategies to Prevent Hospital-onset Staphylococcus aureus Bloodstream Infections in Acute Care Facilities 2024. Available from: https://www.cdc.gov/hai/prevent/staph-prevention-strategies.html.

12. Calderwood MS, Anderson DJ, Bratzler DW, Dellinger EP, Garcia-Houchins S, Maragakis LL, et al. Strategies to prevent surgical site infections in acute-care hospitals: 2022 Update. Infect Control Hosp Epidemiol. 2023;44(5):695–720. Epub 2023/05/04. doi: 10.1017/ice.2023.67. PubMed PMID: 37137483; PubMed Central PMCID: PMCPMC10867741.

13. Popovich KJ, Aureden K, Ham DC, Harris AD, Hessels AJ, Huang SS, et al. SHEA/IDSA/APIC Practice Recommendation: Strategies to prevent methicillin-resistant Staphylococcus aureus transmission and infection in acute-care hospitals: 2022 Update. Infect Control Hosp Epidemiol. 2023;44(7):1039–67. Epub 2023/06/29. doi: 10.1017/ice.2023.102. PubMed PMID: 37381690; PubMed Central PMCID: PMCPMC10369222.

14. Poovelikunnel T, Gethin G, Humphreys H. Mupirocin resistance: clinical implications and potential alternatives for the eradication of MRSA. The Journal of antimicrobial chemotherapy. 2015;70(10):2681–92. Epub 2015/07/05. doi: 10.1093/jac/dkv169. PubMed PMID: 26142407.

15. Poovelikunnel TT, Budri PE, Shore AC, Coleman DC, Humphreys H, Fitzgerald-Hughes D. Molecular Characterization of Nasal Methicillin-Resistant Staphylococcus aureus Isolates Showing Increasing Prevalence of Mupirocin Resistance and Associated Multidrug Resistance following Attempted Decolonization. Antimicrob Agents Chemother. 2018;62(9). Epub 2018/06/20. doi: 10.1128/AAC.00819-18. PubMed PMID: 29914942; PubMed Central PMCID: PMCPMC6125569.

16. Coe KA, Lee W, Stone MC, Komazin-Meredith G, Meredith TC, Grad YH, et al. Multi-strain Tn-Seq reveals common daptomycin resistance determinants in Staphylococcus aureus. PLoS Pathog. 2019;15(11):e1007862. Epub 20191118. doi: 10.1371/journal.ppat.1007862. PubMed PMID: 31738809; PubMed Central PMCID: PMCPMC6934316.

17. Grosser MR, Paluscio E, Thurlow LR, Dillon MM, Cooper VS, Kawula TH, et al. Genetic requirements for Staphylococcus aureus nitric oxide resistance and virulence. PLoS Pathog. 2018;14(3):e1006907. Epub 2018/03/20. doi: 10.1371/journal.ppat.1006907. PubMed PMID: 29554137; PubMed Central PMCID: PMCPMC5884563.

18. Bertrand BP, Shinde D, Thomas VC, Whiteley M, Ibberson CB, Kielian T. Metabolic diversity of human macrophages: potential influence on Staphylococcus aureus intracellular survival. Infect Immun. 2024;92(2):e0047423. Epub 20240105. doi: 10.1128/iai.00474-23. PubMed PMID: 38179975; PubMed Central PMCID: PMCPMC10863412.

19. Chang J, Lee C, Kim I, Kim J, Kim JH, Yun T, et al. Environmental cues in different host niches shape the survival fitness of Staphylococcus aureus. Nat Commun. 2025;16(1):6928. Epub 20250728. doi: 10.1038/s41467-025-62292-x. PubMed PMID: 40721419; PubMed Central PMCID: PMCPMC12304399.

20. Valentino MD, Foulston L, Sadaka A, Kos VN, Villet RA, Santa Maria J, Jr., et al. Genes contributing to Staphylococcus aureus fitness in abscess- and infection-related ecologies. MBio. 2014;5(5):e01729–14. Epub 2014/09/04. doi: 10.1128/mBio.01729-14. PubMed PMID: 25182329; PubMed Central PMCID: PMC4173792.

21. Wilde AD, Snyder DJ, Putnam NE, Valentino MD, Hammer ND, Lonergan ZR, et al. Bacterial Hypoxic Responses Revealed as Critical Determinants of the Host-Pathogen Outcome by TnSeq Analysis of Staphylococcus aureus Invasive Infection. PLoS pathogens. 2015;11(12):e1005341. Epub 2015/12/20. doi: 10.1371/journal.ppat.1005341. PubMed PMID: 26684646; PubMed Central PMCID: PMC4684308.

22. Ibberson CB, Stacy A, Fleming D, Dees JL, Rumbaugh K, Gilmore MS, et al. Co-infecting microorganisms dramatically alter pathogen gene essentiality during polymicrobial infection. Nat Microbiol. 2017;2:17079. Epub 20170530. doi: 10.1038/nmicrobiol.2017.79. PubMed PMID: 28555625; PubMed Central PMCID: PMCPMC5774221.

23. Butrico CE, Klopfenstein N, Green ER, Johnson JR, Peck SH, Ibberson CB, et al. Hyperglycemia Increases Severity of Staphylococcus aureus Osteomyelitis and Influences Bacterial Genes Required for Survival in Bone. Infect Immun. 2023;91(4):e0052922. Epub 20230306. doi: 10.1128/iai.00529-22. PubMed PMID: 36877063; PubMed Central PMCID: PMCPMC10112148.

24. Lyon LM, Marroquin SM, Thorstenson JC, Joyce LR, Fuentes EJ, Doran KS, et al. Genome-wide mutagenesis identifies factors involved in MRSA vaginal olonization. Cell Rep. 2025;44(3):115421. Epub 20250313. doi: 10.1016/j.celrep.2025.115421. PubMed PMID: 40085646; PubMed Central PMCID: PMCPMC12483769.

25. Bilici K, Gerlach D, Camus L, Heilbronner S. Competitive fitness of Staphylococcus aureus against nasal commensals depends on biotin biosynthesis and acquisition. ISME J. 2025;19(1). doi: 10.1093/ismejo/wraf248. PubMed PMID: 41189393; PubMed Central PMCID: PMCPMC12642757.

26. Carfrae LA, MacNair CR, Brown CM, Tsai CN, Weber BS, Zlitni S, et al. Mimicking the human environment in mice reveals that inhibiting biotin biosynthesis is effective against antibiotic-resistant pathogens. Nat Microbiol. 2020;5(1):93–101. Epub 20191028. doi: 10.1038/s41564-019-0595-2. PubMed PMID: 31659298.

27. Yan M, Pamp SJ, Fukuyama J, Hwang PH, Cho DY, Holmes S, et al. Nasal microenvironments and interspecific interactions influence nasal microbiota complexity and *S. aureus* carriage. Cell host & microbe. 2013;14(6):631–40. Epub 2013/12/18. doi: 10.1016/j.chom.2013.11.005. PubMed PMID: 24331461; PubMed Central PMCID: PMC3902146.

28. Rajan A, Weaver AM, Aloisio GM, Jelinski J, Johnson HL, Venable SF, et al. The Human Nose Organoid Respiratory Virus Model: an Ex Vivo Human Challenge Model To Study Respiratory Syncytial Virus (RSV) and Severe Acute Respiratory Syndrome Coronavirus 2 (SARS-CoV-2) Pathogenesis and Evaluate Therapeutics. mBio. 2022;13(1):e0351121. Epub 2022/02/16. doi: 10.1128/mbio.03511-21. PubMed PMID: 35164569; PubMed Central PMCID: PMCPMC8844923.

29. Aloisio GM, Nagaraj D, Murray AM, Schultz EM, McBride T, Aideyan L, et al. Infant-derived human nasal organoids exhibit relatively increased susceptibility, epithelial responses, and cytotoxicity during RSV infection. J Infect. 2024;89(6):106305. Epub 2024/10/11. doi: 10.1016/j.jinf.2024.106305. PubMed PMID: 39389204.

30. Rajan A, Nagaraj D, Bomidi C, Aloisio GM, Murray AM, Schultz EM, et al. Single cell sequencing analysis of respiratory syncytial virus-infected pediatric and adult human nose organoids reveals age differences, proliferative diversity and identifies novel cellular tropism. J Infect. 2025;91(4):106617. Epub 20250918. doi: 10.1016/j.jinf.2025.106617. PubMed PMID: 40975328.

31. Boyd AI, Kafer LA, I FE, Kambal A, Tariq H, Hilsenbeck SG, et al. Nasal microbionts differentially colonize and elicit cytokines in human nasal epithelial organoids. mSphere. 2025;10(10):e0049325. Epub 20250930. doi: 10.1128/msphere.00493-25. PubMed PMID: 41025782; PubMed Central PMCID: PMCPMC12570476.

32. Kafer LA, Escapa IF, Boyd AI, Tostado AR, Kambal A, Blutt SE, et al. Streptococcus pneumoniae colonization modulates human nasal epithelial responses to respiratory syncytial virus infection. bioRxiv. 2026. Epub 20260205. doi: 10.64898/2026.02.05.703985. PubMed PMID: 41676723; PubMed Central PMCID: PMCPMC12889608.

33. Sobral R, Tomasz A. The Staphylococcal Cell Wall. Microbiol Spectr. 2019;7(4). doi: 10.1128/microbiolspec.GPP3-0068-2019. PubMed PMID: 31322105; PubMed Central PMCID: PMCPMC10957225.

34. Peschel A, Otto M, Jack RW, Kalbacher H, Jung G, Gotz F. Inactivation of the dlt operon in Staphylococcus aureus confers sensitivity to defensins, protegrins, and other antimicrobial peptides. J Biol Chem. 1999;274(13):8405–10. doi: 10.1074/jbc.274.13.8405. PubMed PMID: 10085071.

35. Peschel A, Vuong C, Otto M, Gotz F. The D-alanine residues of Staphylococcus aureus teichoic acids alter the susceptibility to vancomycin and the activity of autolytic enzymes. Antimicrob Agents Chemother. 2000;44(10):2845–7. doi: 10.1128/AAC.44.10.2845-2847.2000. PubMed PMID: 10991869; PubMed Central PMCID: PMCPMC90160.

36. Panda S, Jayasinghe YP, Shinde DD, Bueno E, Stastny A, Bertrand BP, et al. Staphylococcus aureus counters organic acid anion-mediated inhibition of peptidoglycan cross-linking through robust alanine racemase activity. eLife. 2024;13:RP95389. Epub 20241106. doi: 10.1101/2024.01.15.575639. PubMed PMID: 38293037; PubMed Central PMCID: PMCPMC10827132.

37. Suzuki Y, Kawada-Matsuo M, Thuan VTT, Le M-T, Sakaguchi T, Komatsuzawa H. D-alanine synthesis and exogenous alanine affect the antimicrobial susceptibility of Staphylococcus aureus. Antimicrob Agents Chemother. 2025;69(7):e0193624. Epub 20250612. doi: 10.1128/aac.01936-24. PubMed PMID: 40503952; PubMed Central PMCID: PMCPMC12217452.

38. Anthony KG, Strych U, Yeung KR, Shoen CS, Perez O, Krause KL, et al. New classes of alanine racemase inhibitors identified by high-throughput screening show antimicrobial activity against Mycobacterium tuberculosis. PLoS One. 2011;6(5):e20374. Epub 20110526. doi: 10.1371/journal.pone.0020374. PubMed PMID: 21637807; PubMed Central PMCID: PMCPMC3102704.

39. Wei Y, Qiu W, Zhou XD, Zheng X, Zhang KK, Wang SD, et al. Alanine racemase is essential for the growth and interspecies competitiveness of Streptococcus mutans. Int J Oral Sci. 2016;8(4):231–8. Epub 20161216. doi: 10.1038/ijos.2016.34. PubMed PMID: 27740612; PubMed Central PMCID: PMCPMC5168415.

40. de Chiara C, Homsak M, Prosser GA, Douglas HL, Garza-Garcia A, Kelly G, et al. D-Cycloserine destruction by alanine racemase and the limit of irreversible inhibition. Nature chemical biology. 2020;16(6):686–94. Epub 20200316. doi: 10.1038/s41589-020-0498-9. PubMed PMID: 32203411; PubMed Central PMCID: PMCPMC7246083.

41. Moscoso M, Garcia P, Cabral MP, Rumbo C, Bou G. A D-Alanine auxotrophic live vaccine is effective against lethal infection caused by Staphylococcus aureus. Virulence. 2018;9(1):604–20. doi: 10.1080/21505594.2017.1417723. PubMed PMID: 29297750; PubMed Central PMCID: PMCPMC5955480.

42. Mulcahy ME, McLoughlin RM. Host-Bacterial Crosstalk Determines Staphylococcus aureus Nasal Colonization. Trends Microbiol. 2016;24(11):872–86. Epub 20160726. doi: 10.1016/j.tim.2016.06.012. PubMed PMID: 27474529.

43. Lin MH, Shu JC, Lin LP, Chong KY, Cheng YW, Du JF, et al. Elucidating the crucial role of poly N-acetylglucosamine from Staphylococcus aureus in cellular adhesion and pathogenesis. PLoS One. 2015;10(4):e0124216. Epub 20150415. doi: 10.1371/journal.pone.0124216. PubMed PMID: 25876106; PubMed Central PMCID: PMCPMC4398431.

44. Huffines JT, Boone RL, Kiedrowski MR. Temperature influences commensal-pathogen dynamics i a nasal epithelial cell co-culture model. mSphere. 2024;9(1):e0058923. Epub 2024/01/05. doi: 10.1128/msphere.00589-23. PubMed PMID: 38179905; PubMed Central PMCID: PMCPMC10826359.

45. Huffines JT, Kiedrowski MR. Staphylococcus aureus phenol-soluble modulins have dispersal and anti-aggregation activity towards corynebacteria. J Bacteriol. 2025;207(9):e0018325. Epub 20250814. doi: 10.1128/jb.00183-25. PubMed PMID: 40810517; PubMed Central PMCID: PMCPMC12445096.

46. Miller LG, Perdreau-Remington F, Rieg G, Mehdi S, Perlroth J, Bayer AS, et al. Necrotizing fasciitis caused by community-associated methicillin-resistant Staphylococcus aureus in Los Angeles. N Engl J Med. 2005;352(14):1445–53. PubMed PMID: 15814880.

47. Keck T, Leiacker R, Riechelmann H, Rettinger G. Temperature profile in the nasal cavity. Laryngoscope. 2000;110(4):651–4. Epub 2000/04/14. doi: 10.1097/00005537-200004000-00021. PubMed PMID: 10764013.

48. Bastock RA, Marino EC, Wiemels RE, Holzschu DL, Keogh RA, Zapf RL, et al. Staphylococcus aureus Responds to Physiologically Relevant Temperature Changes by Altering Its Global Transcript and Protein Profile. mSphere. 2021;6(2). Epub 2021/03/19. doi: 10.1128/mSphere.01303-20. PubMed PMID: 33731473; PubMed Central PMCID: PMCPMC8546721.

49. Kiedrowski MR, Paharik AE, Ackermann LW, Shelton AU, Singh SB, Starner TD, et al. Development of an in vitro colonization model to investigate *Staphylococcus aureus* interactions with airway epithelia. Cell Microbiol. 2016;18(5):720–32. Epub 2015/11/14. doi: 10.1111/cmi.12543. PubMed PMID: 26566259; PubMed Central PMCID: PMCPMC4840028.

50. Turner KH, Everett J, Trivedi U, Rumbaugh KP, Whiteley M. Requirements for Pseudomonas aeruginosa acute burn and chronic surgical wound infection. PLoS genetics. 2014;10(7):e1004518. Epub 2014/07/25. doi: 10.1371/journal.pgen.1004518. PubMed PMID: 25057820; PubMed Central PMCID: PMC4109851.

51. Stacy A, Fleming D, Lamont RJ, Rumbaugh KP, Whiteley M. A Commensal Bacterium Promotes Virulence of an Opportunistic Pathogen via Cross-Respiration. MBio. 2016;7(3). Epub 2016/06/30. doi: 10.1128/mBio.00782-16. PubMed PMID: 27353758; PubMed Central PMCID: PMC4916382.

52. Lewin GR, Stacy A, Michie KL, Lamont RJ, Whiteley M. Large-scale identification of pathogen essential genes during coinfection with sympatric and allopatric microbes. Proc Natl Acad Sci U S A. 2019;116(39):19685–94. Epub 20190819. doi: 10.1073/pnas.1907619116. PubMed PMID: 31427504; PubMed Central PMCID: PMCPMC6765283.

53. The Gene Ontology C. The Gene Ontology Resource: 20 years and still GOing strong. Nucleic Acids Res. 2019;47(D1):D330–D8. doi: 10.1093/nar/gky1055. PubMed PMID: 30395331; PubMed Central PMCID: PMCPMC6323945.

54. Szklarczyk D, Kirsch R, Koutrouli M, Nastou K, Mehryary F, Hachilif R, et al. The STRING database in 2023: protein-protein association networks and functional enrichment analyses for any sequenced genome of interest. Nucleic Acids Res. 2023;51(D1):D638–D46. doi: 10.1093/nar/gkac1000. PubMed PMID: 36370105; PubMed Central PMCID: PMCPMC9825434.

55. Snel B, Lehmann G, Bork P, Huynen MA. STRING: a web-server to retrieve and display the repeatedly occurring neighbourhood of a gene. Nucleic Acids Res. 2000;28(18):3442–4. doi: 10.1093/nar/28.18.3442. PubMed PMID: 10982861; PubMed Central PMCID: PMCPMC110752.

56. Grim KP, San Francisco B, Radin JN, Brazel EB, Kelliher JL, Parraga Solorzano PK, et al. The Metallophore Staphylopine Enables Staphylococcus aureus To Compete with the Host for Zinc and Overcome Nutritional Immunity. mBio. 2017;8(5). Epub 20171031. doi: 10.1128/mBio.01281-17. PubMed PMID: 29089427; PubMed Central PMCID: PMCPMC5666155.

57. Bulock LL, Ahn J, Shinde D, Pandey S, Sarmiento C, Thomas VC, et al. Interplay of CodY and CcpA in Regulating Central Metabolism and Biofilm Formation in Staphylococcus aureus. J Bacteriol. 2022;204(7):e0061721. Epub 20220623. doi: 10.1128/jb.00617-21. PubMed PMID: 35735992; PubMed Central PMCID: PMCPMC9295537.

58. Nuxoll AS, Halouska SM, Sadykov MR, Hanke ML, Bayles KW, Kielian T, et al. CcpA regulates arginine biosynthesis in Staphylococcus aureus through repression of proline catabolism. PLoS Pathog. 2012;8(11):e1003033. Epub 20121129. doi: 10.1371/journal.ppat.1003033. PubMed PMID: 23209408; PubMed Central PMCID: PMCPMC3510247.

59. Goncheva MI, Flannagan RS, Heinrichs DE. De Novo Purine Biosynthesis Is Required for Intracellular Growth of Staphylococcus aureus and for the Hypervirulence Phenotype of a purR Mutant. Infect Immun. 2020;88(5). Epub 20200420. doi: 10.1128/IAI.00104-20. PubMed PMID: 32094249; PubMed Central PMCID: PMCPMC7171247.

60. Kofoed EM, Yan D, Katakam AK, Reichelt M, Lin B, Kim J, et al. De Novo Guanine Biosynthesis but Not the Riboswitch-Regulated Purine Salvage Pathway Is Required for Staphylococcus aureus Infection In Vivo. J Bacteriol. 2016;198(14):2001–15. Epub 20160627. doi: 10.1128/JB.00051-16. PubMed PMID: 27161118; PubMed Central PMCID: PMCPMC4936099.

61. Mulhbacher J, Brouillette E, Allard M, Fortier LC, Malouin F, Lafontaine DA. Novel riboswitch ligand analogs as selective inhibitors of guanine-related metabolic pathways. PLoS Pathog. 2010;6(4):e1000865. Epub 20100422. doi: 10.1371/journal.ppat.1000865. PubMed PMID: 20421948; PubMed Central PMCID: PMCPMC2858708.

62. Li L, Abdelhady W, Donegan NP, Seidl K, Cheung A, Zhou YF, et al. Role of Purine Biosynthesis in Persistent Methicillin-Resistant Staphylococcus aureus Infection. J Infect Dis. 2018;218(9):1367–77. doi: 10.1093/infdis/jiy340. PubMed PMID: 29868791; PubMed Central PMCID: PMCPMC6151072.

63. Berry KA, Verhoef MTA, Zheng Z, Flannagan RS, Paiva TO, Gilbert SE, et al. The Staphylococcus aureus esterase FmtA is essential for wall teichoic acid D-alanylation. mBio. 2025;16(10):e0233725. Epub 20250912. doi: 10.1128/mbio.02337-25. PubMed PMID: 40937855; PubMed Central PMCID: PMCPMC12505958.

64. Schultz BJ, Snow ED, Walker S. Mechanism of D-alanine transfer to teichoic acids shows how bacteria acylate cell envelope polymers. Nat Microbiol. 2023;8(7):1318–29. Epub 20230612. doi: 10.1038/s41564-023-01411-0. PubMed PMID: 37308592; PubMed Central PMCID: PMCPMC10664464.

65. Fisher SL. Glutamate racemase as a target for drug discovery. Microb Biotechnol. 2008;1(5):345–60. Epub 20080511. doi: 10.1111/j.1751-7915.2008.00031.x. PubMed PMID: 21261855; PubMed Central PMCID: PMCPMC3815242.

66. Bearne SL. Through the Looking Glass: Chiral Recognition of Substrates and Products at the Active Sites of Racemases and Epimerases. Chemistry (Weinheim an der Bergstrasse, Germany). 2020;26(46):10367–90. Epub 20200721. doi: 10.1002/chem.201905826. PubMed PMID: 32166792.

67. Chaudhuri RR, Allen AG, Owen PJ, Shalom G, Stone K, Harrison M, et al. Comprehensive identification of essential Staphylococcus aureus genes using Transposon-Mediated Differential Hybridisation (TMDH). BMC Genomics. 2009;10:291. Epub 20090701. doi: 10.1186/1471-2164-10-291. PubMed PMID: 19570206; PubMed Central PMCID: PMCPMC2721850.

68. Luong TT, Lee CY. Improved single-copy integration vectors for Staphylococcus aureus. J Microbiol Methods. 2007;70(1):186–90. Epub 20070424. doi: 10.1016/j.mimet.2007.04.007. PubMed PMID: 17512993; PubMed Central PMCID: PMCPMC2001203.

69. Burlak C, Hammer CH, Robinson MA, Whitney AR, McGavin MJ, Kreiswirth BN, et al. Global analysis of community-associated methicillin-resistant Staphylococcus aureus exoproteins reveals molecules produced in vitro and during infection. Cell Microbiol. 2007;9(5):1172–90. Epub 20070109. doi: 10.1111/j.1462-5822.2006.00858.x. PubMed PMID: 17217429; PubMed Central PMCID: PMCPMC2064037.

70. Goldstein JM, Nelson D, Kordula T, Mayo JA, Travis J. Extracellular arginine aminopeptidase from Streptococcus gordonii FSS2. Infect Immun. 2002;70(2):836–43. doi: 10.1128/IAI.70.2.836-843.2002. PubMed PMID: 11796618; PubMed Central PMCID: PMCPMC127726.

71. Coppens L, Lavigne R. SAPPHIRE: a neural network based classifier for sigma70 promoter prediction in Pseudomonas. BMC Bioinformatics. 2020;21(1):415. Epub 20200922. doi: 10.1186/s12859-020-03730-z. PubMed PMID: 32962628; PubMed Central PMCID: PMCPMC7510298.

72. Tran TH, Escapa IF, Roberts AQ, Gao W, Obawemimo AC, Segre JA, et al. Metabolic capabilities are highly conserved among human nasal-associated Corynebacterium species in pangenomic analyses. mSystems. 2024;9(12):e0113224. Epub 2024/11/13. doi: 10.1128/msystems.01132-24. PubMed PMID: 39508593; PubMed Central PMCID: PMCPMC11651106.

73. Krismer B, Liebeke M, Janek D, Nega M, Rautenberg M, Hornig G, et al. Nutrient limitation governs *Staphylococcus aureus* metabolism and niche adaptation in the human nose. PLoS Pathog. 2014;10(1):e1003862. Epub 2014/01/24. doi: 10.1371/journal.ppat.1003862 PPATHOGENS-D-13-02760 [pii]. PubMed PMID: 24453967; PubMed Central PMCID: PMCPMC3894218.

74. Farne H, Groves HT, Gill SK, Stokes I, McCulloch S, Karoly E, et al. Comparative Metabolomic Sampling of Upper and Lower Airways by Four Different Methods to Identify Biochemicals That May Support Bacterial Growth. Front Cell Infect Microbiol. 2018;8:432. Epub 2019/01/09. doi: 10.3389/fcimb.2018.00432. PubMed PMID: 30619778; PubMed Central PMCID: PMCPMC6305596.

75. Figueiredo TA, Sobral RG, Ludovice AM, Almeida JM, Bui NK, Vollmer W, et al. Identification of genetic determinants and enzymes involved with the amidation of glutamic acid residues in the peptidoglycan of Staphylococcus aureus. PLoS Pathog. 2012;8(1):e1002508. Epub 20120126. doi: 10.1371/journal.ppat.1002508. PubMed PMID: 22303291; PubMed Central PMCID: PMCPMC3267633.

76. Fey PD, Endres JL, Yajjala VK, Widhelm TJ, Boissy RJ, Bose JL, et al. A genetic resource for rapid and comprehensive phenotype screening of nonessential *Staphylococcus aureus* genes. mBio. 2013;4(1):e00537–12. Epub 2013/02/14. doi: 10.1128/mBio.00537-12. PubMed PMID: 23404398; PubMed Central PMCID: PMCPMC3573662.

77. Shinefield HR, Ribble JC, Boris M, Eichenwald HF. Bacterial interference: its effect on nursery-acquired infection with Staphylococcus aureus. I. Preliminary observations on artificial colonzation of newborns. Am J Dis Child. 1963;105:646–54. PubMed PMID: 13977323.

78. Parker D, Narechania A, Sebra R, Deikus G, Larussa S, Ryan C, et al. Genome Sequence of Bacterial Interference Strain Staphylococcus aureus 502A. Genome Announc. 2014;2(2). Epub 2014/04/12. doi: 10.1128/genomeA.00284-14. PubMed PMID: 24723721; PubMed Central PMCID: PMCPMC3983310.

79. Chapman JR, Balasubramanian D, Tam K, Askenazi M, Copin R, Shopsin B, et al. Using Quantitative Spectrometry to Understand the Influence of Genetics and Nutritional Perturbations On the Virulence Potential of Staphylococcus aureus. Mol Cell Proteomics. 2017;16(4 suppl 1):S15-S28. Epub 20170214. doi: 10.1074/mcp.O116.065581. PubMed PMID: 28196877; PubMed Central PMCID: PMCPMC5393389.

80. Mootz JM, Benson MA, Heim CE, Crosby HA, Kavanaugh JS, Dunman PM, et al. Rot is a key regulator of Staphylococcus aureus biofilm formation. Molecular microbiology. 2015;96(2):388–404. Epub 20150226. doi: 10.1111/mmi.12943. PubMed PMID: 25612137; PubMed Central PMCID: PMCPMC4467170.

81. Reichmann NT, Cassona CP, Grundling A. Revised mechanism of D-alanine incorporation into cell wall polymers in Gram-positive bacteria. Microbiology (Reading). 2013;159(Pt 9):1868–77. Epub 20130715. doi: 10.1099/mic.0.069898-0. PubMed PMID: 23858088; PubMed Central PMCID: PMCPMC3783018.

82. Collins LV, Kristian SA, Weidenmaier C, Faigle M, Van Kessel KP, Van Strijp JA, et al. Staphylococcus aureus strains lacking D-alanine modifications of teichoic acids are highly susceptible to human neutrophil killing and are virulence attenuated in mice. J Infect Dis. 2002;186(2):214–9. Epub 20020703. doi: 10.1086/341454. PubMed PMID: 12134257.

83. Brugger SD, Eslami SM, Pettigrew MM, Escapa IF, Henke MT, Kong Y, et al. *Dolosigranulum pigrum* Cooperation and Competition in Human Nasal Microbiota. mSphere. 2020;5(5):e00852–20. Epub 2020/09/11. doi: 10.1128/mSphere.00852-20. PubMed PMID: 32907957; PubMed Central PMCID: PMCPMC7485692.

84. Escapa IF, Chen T, Huang Y, Gajare P, Dewhirst FE, Lemon KP. New Insights into Human Nostril Microbiome from the Expanded Human Oral Microbiome Database (eHOMD): a Resource for the Microbiome of the Human Aerodigestive Tract. mSystems. 2018;3(6):e00187–18. Epub 2018/12/12. doi: 10.1128/mSystems.00187-18. PubMed PMID: 30534599; PubMed Central PMCID: PMCPMC6280432.

85. Roy R, Jayasinghe YP, Panda S, Zeden MS, Thomas VC, Ronning DR, et al. A S(180)F substitution in D-alanine aminotransferase confers resistance to beta-chloro-D-alanine in Staphylococcus aureus. J Biol Chem. 2025;301(12):110931. Epub 20251111. doi: 10.1016/j.jbc.2025.110931. PubMed PMID: 41232672; PubMed Central PMCID: PMCPMC12723384.

86. Lensmire JM, Wischer MR, Kraemer-Zimpel C, Kies PJ, Sosinski L, Ensink E, et al. The glutathione import system satisfies the Staphylococcus aureus nutrient sulfur requirement and promotes interspecies competition. PLoS Genet. 2023;19(7):e1010834. Epub 20230707. doi: 10.1371/journal.pgen.1010834. PubMed PMID: 37418503; PubMed Central PMCID: PMCPMC10355420.

87. Hiron A, Borezee-Durant E, Piard JC, Juillard V. Only one of four oligopeptide transport systems mediates nitrogen nutrition in Staphylococcus aureus. J Bacteriol. 2007;189(14):5119–29. Epub 20070511. doi: 10.1128/JB.00274-07. PubMed PMID: 17496096; PubMed Central PMCID: PMCPMC1951871.

88. Shirai T, Takase D, Yokoyama J, Nakanishi K, Uehara C, Saito N, et al. Functions of human olfactory mucus and age-dependent changes. Sci Rep. 2023;13(1):971. Epub 20230118. doi: 10.1038/s41598-023-27937-1. PubMed PMID: 36653421; PubMed Central PMCID: PMCPMC9846672.

89. Halsey CR, Lei S, Wax JK, Lehman MK, Nuxoll AS, Steinke L, et al. Amino Acid Catabolism in Staphylococcus aureus and the Function of Carbon Catabolite Repression. mBio. 2017;8(1). Epub 20170214. doi: 10.1128/mBio.01434-16. PubMed PMID: 28196956; PubMed Central PMCID: PMCPMC5312079.

90. Wood DM, Brennan AL, Philips BJ, Baker EH. Effect of hyperglycaemia on glucose concentration of human nasal secretions. Clin Sci (Lond). 2004;106(5):527–33. doi: 10.1042/CS20030333. PubMed PMID: 14678009.

91. Cabral MP, Garcia P, Beceiro A, Rumbo C, Perez A, Moscoso M, et al. Design of live attenuated bacterial vaccines based on D-glutamate auxotrophy. Nat Commun. 2017;8:15480. Epub 20170526. doi: 10.1038/ncomms15480. PubMed PMID: 28548079; PubMed Central PMCID: PMCPMC5458566.

92. Sidiq KR, Chow MW, Zhao Z, Daniel RA. Alanine metabolism in Bacillus subtilis. Molecular microbiology. 2021;115(4):739–57. Epub 20201124. doi: 10.1111/mmi.14640. PubMed PMID: 33155333.

93. Koper K, Han SW, Kothadia R, Salamon H, Yoshikuni Y, Maeda HA. Multisubstrate specificity shaped the complex evolution of the aminotransferase family across the tree of life. Proc Natl Acad Sci U S A. 2024;121(26):e2405524121. Epub 20240617. doi: 10.1073/pnas.2405524121. PubMed PMID: 38885378; PubMed Central PMCID: PMCPMC11214133.

94. Flores Ramos S, Brugger SD, Escapa IF, Skeete CA, Cotton SL, Eslami SM, et al. Genomic Stability and Genetic Defense Systems in *Dolosigranulum pigrum*, a Candidate Beneficial Bacterium from the Human Microbiome. mSystems. 2021;6(5):e0042521. Epub 2021/09/22. doi: 10.1128/mSystems.00425-21. PubMed PMID: 34546072; PubMed Central PMCID: PMCPMC8547433.

95. Bose JL, Fey PD, Bayles KW. Genetic tools to enhance the study of gene function and regulation in Staphylococcus aureus. Appl Environ Microbiol. 2013;79(7):2218–24. Epub 2013/01/29. doi: 10.1128/AEM.00136-13 AEM.00136-13 [pii]. PubMed PMID: 23354696; PubMed Central PMCID: PMC3623228.

96. Klein BA, Tenorio EL, Lazinski DW, Camilli A, Duncan MJ, Hu LT. Identification of essential genes of the periodontal pathogen Porphyromonas gingivalis. BMC Genomics. 2012;13:578. Epub 2012/11/02. doi: 10.1186/1471-2164-13-578. PubMed PMID: 23114059; PubMed Central PMCID: PMC3547785.

97. Martin M. Cutadapt Removes Adapter Sequences from High-Throughput Sequencing Reads. EMBnet Journal. 2011;17:10–2. doi: 10.14806/ej.17.1.200.

98. Langmead B, Salzberg SL. Fast gapped-read alignment with Bowtie 2. Nat Methods. 2012;9(4):357–9. Epub 20120304. doi: 10.1038/nmeth.1923. PubMed PMID: 22388286; PubMed Central PMCID: PMCPMC3322381.

99. Langmead B, Wilks C, Antonescu V, Charles R. Scaling read aligners to hundreds of threads on general-purpose processors. Bioinformatics. 2019;35(3):421–32. doi: 10.1093/bioinformatics/bty648. PubMed PMID: 30020410; PubMed Central PMCID: PMCPMC6361242.

100. Danecek P, Bonfield JK, Liddle J, Marshall J, Ohan V, Pollard MO, et al. Twelve years of SAMtools and BCFtools. GigaScience. 2021;10(2). doi: 10.1093/gigascience/giab008. PubMed PMID: 33590861; PubMed Central PMCID: PMCPMC7931819.

101. Love MI, Huber W, Anders S. Moderated estimation of fold change and dispersion for RNA-seq data with DESeq2. Genome Biol. 2014;15(12):550. Epub 2014/12/18. doi: 10.1186/s13059-014-0550-8. PubMed PMID: 25516281; PubMed Central PMCID: PMCPMC4302049.

102. Scrucca L, Fraley C, Murphy TB, Raftery AE. Model-Based Clustering, Classification, and Density Estimation Using mclust in R: Chapman and Hall/CRC.; 2023.

103. Zeden MS, Schuster CF, Grundling A. Preparation of Electrocompetent Staphylococcus aureus Cells and Plasmid Transformation. Cold Spring Harb Protoc. 2023;2023(8):107947. Epub 20230801. doi: 10.1101/pdb.prot107947. PubMed PMID: 37117021.

104. Kreiswirth BN, Lofdahl S, Betley MJ, O’Reilly M, Schlievert PM, Bergdoll MS, et al. The toxic shock syndrome exotoxin structural gene is not detectably transmitted by a prophage. Nature. 1983;305(5936):709–12. doi: 10.1038/305709a0. PubMed PMID: 6226876.

105. Duran Ramirez JM, Gomez J, Obernuefemann CLP, Gualberto NC, Walker JN. Semi-Quantitative Assay to Measure Urease Activity by Urinary Catheter-Associated Uropathogens. Front Cell Infect Microbiol. 2022;12:859093. Epub 20220322. doi: 10.3389/fcimb.2022.859093. PubMed PMID: 35392611; PubMed Central PMCID: PMCPMC8980526.

106. Dower WJ, Miller JF, Ragsdale CW. High efficiency transformation of E. coli by high voltage electroporation. Nucleic Acids Res. 1988;16(13):6127–45. doi: 10.1093/nar/16.13.6127. PubMed PMID: 3041370; PubMed Central PMCID: PMCPMC336852.

107. R-Core-Team. R: A Language and Environment for Statistical Computing. Vienna, Austria: R Foundation for Statistical Computing; 2021.

108. RStudio-Team. RStudio: Integrated Development for R. RStudio. Boston, MA: PBC; 2020.

109. Ushey K, Wickham H. renv: Project Environments. R package version 1.2.3 ed2026.

110. Bates D, Mächler M, Bolker B, Walker S. Fitting Linear Mixed-Effects Models Using lme4. Journal of Statistical Software. 2015;67(1):1–48. doi: 10.18637/jss.v067.i01.

111. Kuznetsova A, Brockhoff BP, Christensen BHR. lmerTest Package: Tests in Linear Mixed Effects Models. Journal of Statistical Software. 2017;82(13). doi: doi.org/10.18637/jss.v082.i13.

112. Lenth RV. emmeans: Estimated Marginal Means, aka Least-Squares Means. 2024.

113. Holm S. A Simple Sequentially Rejective Multiple Test Procedure. Scandinavian Journal of Statistics. 1979;6(2):65–70.

